# Structure of the γ-tubulin ring complex-capped microtubule

**DOI:** 10.1101/2023.11.20.567916

**Authors:** Amol Aher, Linas Urnavicius, Allen Xue, Kasahun Neselu, Tarun M. Kapoor

## Abstract

Microtubules are composed of α/β-tubulin dimers positioned head-to-tail to form protofilaments that associate laterally in varying numbers. It is not known how cellular microtubules assemble with the canonical 13-protofilament architecture, resulting in micrometer-scale α/β-tubulin tracks for intracellular transport that align with, rather than spiral along, the filament’s long-axis. We report that the human ∼2.3MDa γ-tubulin ring complex (γ-TuRC), an essential regulator of microtubule formation that contains 14 γ-tubulins, selectively nucleates 13-protofilament microtubules. Cryo-EM reconstructions of γ-TuRC-capped microtubule minus-ends reveal the extensive intra- and inter-domain motions of γ-TuRC subunits that accommodate its actin-containing luminal bridge and establish lateral and longitudinal interactions between γ- and α-tubulins. Our structures reveal how free γ-TuRC, an inefficient nucleation template due to its splayed conformation, transforms into a stable cap that blocks addition or loss of α/β-tubulins from minus-ends and sets the lattice architecture of cellular microtubules.

**One Sentence Summary:** Structural insights into how the γ-tubulin ring complex nucleates and caps a 13-protofilament microtubule.

## INTRODUCTION

Microtubules are cytoskeletal polymers required for essential cellular processes such as directional transport and cell division (*1–3*). Since the 1970s, it has been known that in eukaryotic cells microtubules are typically composed of 13 protofilaments, which are formed by α/β-tubulin dimers that align longitudinally (*4*). In contrast, microtubules assembled *in vitro*, with purified α/β-tubulin, have 8-16 protofilaments (*5–7*). The γ-tubulin ring complex (γ-TuRC), an essential microtubule nucleator that has 14 γ-tubulins, GCP2-GCP6 (γ-tubulin complex proteins-2 to -6) and an actin-containing luminal bridge assembled into a cone-shaped structure (*8–12*), could provide a template that sets the lattice architecture of cellular microtubules (*13*). However, it is unclear if γ-TuRC can nucleate 13-protofilament microtubules.

Studies of preparations of *Drosophila* γ-TuRCs and centrosomes, yeast spindle pole bodies and *Xenopus* extracts along with immunogold-based analyses, have provided evidence for microtubule minus-ends capped with discernible cone-shaped densities and support a ‘template’ model for γ-TuRC function (*14–17*). In this model, γ-tubulins in the γ-TuRCs directly bind α-tubulins at the microtubule minus ends and align with each protofilament longitudinally (*18*, *19*). The yeast γ-TuSC (γ-tubulin small complex) is a V-shaped complex consisting of only two γ-tubulins, GCP2 and GCP3 (*20–22*), and under specific conditions can assemble into extended spiral structures *in vitro* (*23*). The positions of γ-tubulins in these assemblies do not properly match, without engineered disulfide bonds, the positions of α/β-tubulin in the canonical microtubule lattice (*17*). Recent cryo-electron microscopy (cryo-EM) structures of human and *Xenopus* γ-TuRC have revealed that GCP2-GCP6 are arranged into 14 spokes that position 14 γ-tubulins in a lock-washer shape (*8–11*). The even number of γ-tubulins in these complexes, similar to the yeast γ-TuSC assemblies, are found in orientations and positions (e.g. helical rise and pitch) that do not match the geometry needed to bind the odd number of α-tubulins at the minus ends of a 13-protofilament microtubule (*8*, *9*, *23*). Consistent with these structural data, the efficiency of microtubule nucleation by γ-TuRC *in vitro* is low (∼0.5%) (*10, 24*). In contrast, γ-TuRC caps the minus-ends of nucleated microtubules with long lifetimes (10-30 minutes) (*25*), suggesting that the complex can adopt a shape that matches the microtubule minus-end. However, structural evidence for γ-TuRC-dependent templated microtubule nucleation is lacking and it is unclear how the free γ-TuRC transforms into a stable minus-end cap.

To address these open questions we used cryo-EM to show that the γ-TuRC selectively nucleates 13-protofilament microtubules and determined the structure of a γ-TuRC-capped microtubule minus-end.

## RESULTS

### The γ-TuRC selectively nucleates 13-protofilament microtubules

To examine the protofilament number of microtubules nucleated by γ-TuRC we optimized a nucleation assay on cryo-EM grids using bovine tubulin and native human γ-TuRC purified from HeLa S3 cells (materials and methods; Fig. 1A and fig. S1A,B). While 1000s of γ-TuRC complexes (∼30 nm in diameter) could be readily concentrated in a ‘hole’ on the grid, as has been done for single-particle studies (*8–10*), it was difficult to get sufficient numbers of γ-TuRC-capped microtubule ends, in large part due to the micrometer lengths of microtubules and γ-TuRC’s low nucleation efficiency (∼0.5%) (*10*, *24*). Inclusion of chTOG, a polymerase (*26*), increased the nucleation rate (*10*, *27*), and we were able to collect cryo-EM images to identify 177 microtubules capped by a cone-shaped density at their ends (materials and methods; Fig. 1B, white arrows). Consistent with earlier studies (*28*, *29*), analysis of spontaneously nucleated microtubules revealed that 46.8 ± 6.1 % had 13 protofilaments and 51.1 ± 6.9% had 14 protofilaments (Fig. 1C; materials and methods). In contrast, 92 ± 7.1% of microtubules nucleated by native γ-TuRC had 13 protofilaments (Fig. 1D), matching the canonical architecture of cellular microtubules (*30*).

**Fig. 1.**
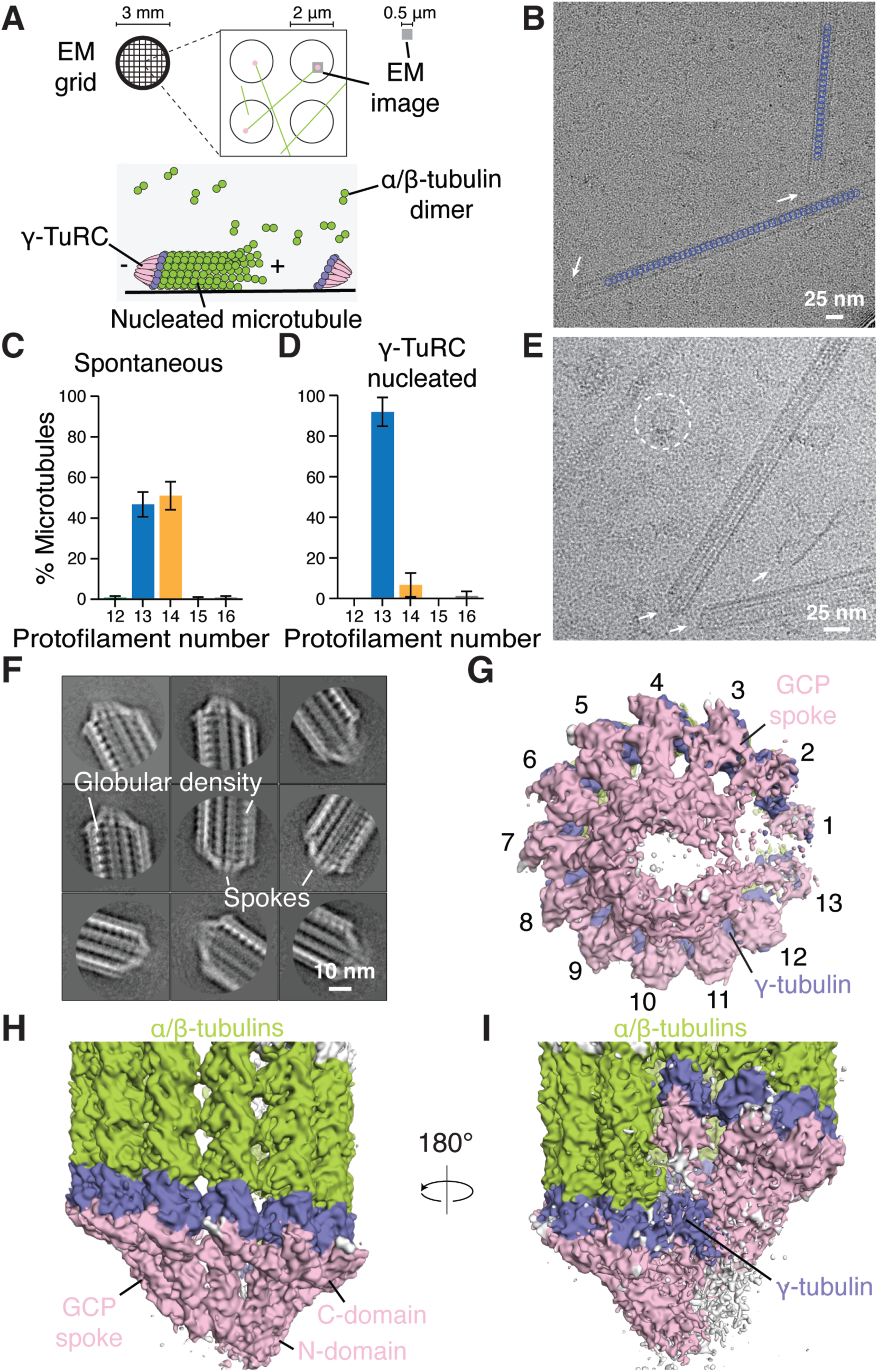
Cryo-EM of γ-TuRC-capped microtubules reveals selective nucleation of 13-protofilament microtubules. (**A**) Schematic of the on-grid nucleation assay for cryo-EM-based analysis. (**B**) Micrograph showing two microtubules nucleated in the presence of native human γ-TuRC. Cone-shaped densities at microtubule ends, consistent with γ-TuRC caps, are indicated (white arrow). Segment assignments by 3D classification indicate protofilament number (blue circles: 13). (**C, D**) Percent microtubules with different protofilament numbers nucleated in the absence (C) or presence (D) of native γ-TuRC. (**E**) Micrograph showing microtubules with capped ends (arrows) and free recombinant γ-TuRC (circle). (**F**) 2D class averages of capped microtubule ends reveal repeating globular and spoke-like densities. (**G-I**) Surface representation of the overall density for γ-TuRC-capped microtubule end in different views (colored as indicated).

For structural studies of γ-TuRC-capped microtubules, we switched to using recombinant γ-TuRC (fig. S1C) (*24*), which could be purified to concentrations higher than the native complex, and recombinant isotypically-pure human tubulin with an ɑ-tubulin mutation (E254D) that slows GTP hydrolysis (*31*). Optimized conditions yielded 92 ± 3.0% of microtubules nucleated by γ-TuRC (fig. S1D-G). Following synchronous and rapid (∼50 s) on-grid nucleation, we obtained micrographs in which free γ-TuRC particles (Fig. 1E, circle) and microtubules with a cone-shaped density at one end (Fig. 1E, white arrows) could be identified. From ∼75,000 micrographs we picked ∼45,000 capped microtubules (fig. S2).

Reference-free 2D classification of capped microtubule ends showed repeating globular densities that were consistent with α/β-tubulins arranged into protofilaments (Fig. 1F). Importantly, we also observed ‘spokes’ within the cone-shaped density at microtubule ends (Fig. 1F). After several rounds of reference-free 2D classification, we selected 24,406 clean ‘ends’ that were used for a 3D reconstruction (7.7Å resolution) (Fig. 1G and fig. S2). At least 13 individual spokes, consistent with GCP domains (pink), organized into a cone (Fig. 1G) were apparent. Distal to the vertex of the cone, each GCP density was followed by a globular density consistent with γ-tubulin (blue), and additional globular densities consistent with α/β-tubulin subunits (green) (Fig. 1H,I). This reconstruction reveals the overall organization of a γ-TuRC-bound to the minus-end of a 13-protofilament microtubule, which we find this complex selectively nucleates.

### Longitudinal interactions at the γ-TuRC capped microtubule minus-end

Crosssectional view of the filament interior revealed a density consistent with the ‘luminal bridge’ (orange, Fig. 2A) of the γ-TuRC (*8–11*). In addition, stronger densities were observed for 10 protofilaments and the associated GCPs, while the other densities were weaker, consistent with higher conformational dynamics and/or heterogeneity in this region (Fig. 2A). To analyze the longitudinal interface between γ-TuRC and the microtubule minus-end we focused on the stable side (positions 3-9) of the γ-TuRC-capped microtubule to improve resolution.

**Fig. 2.**
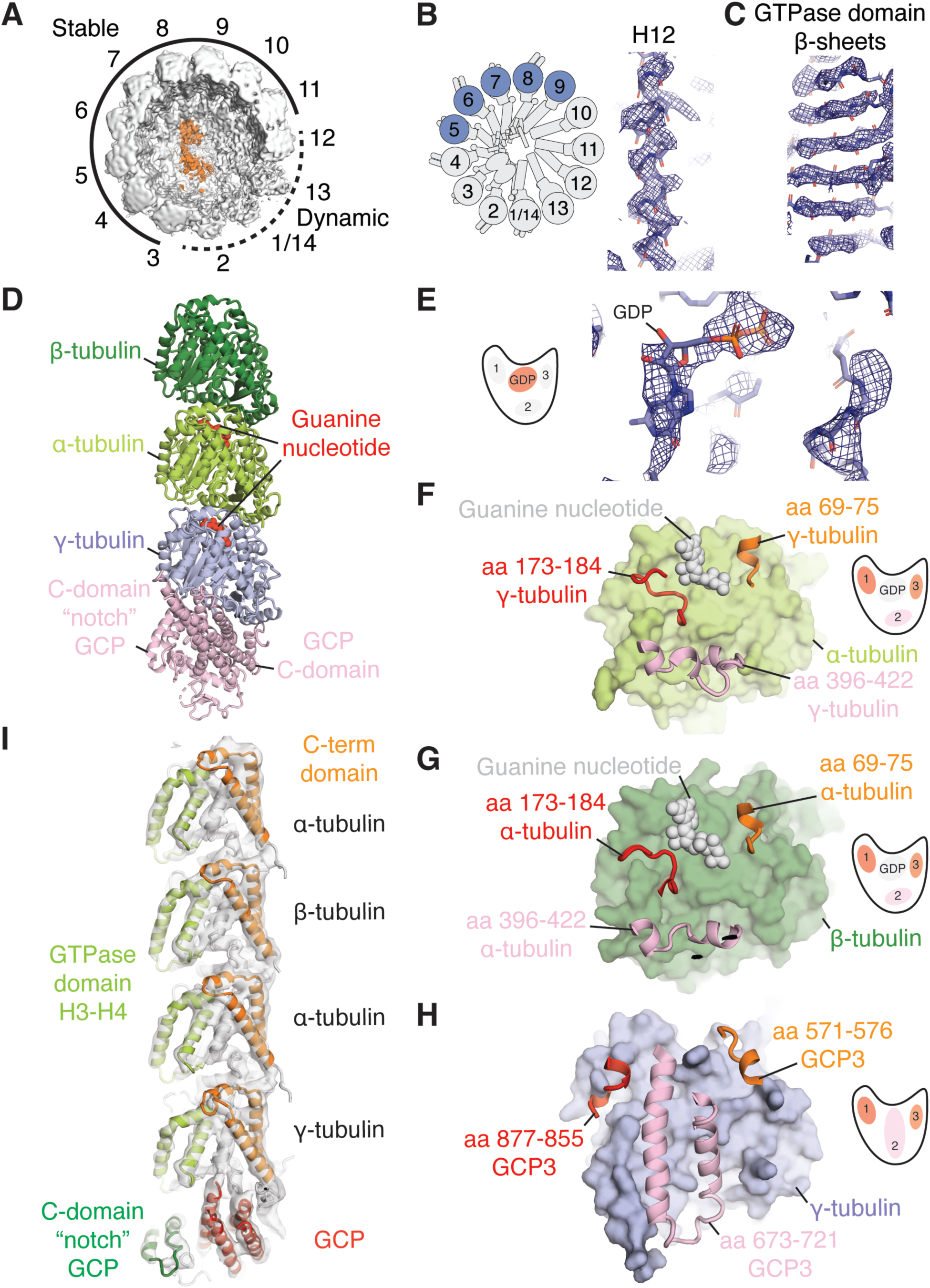
Longitudinal interactions along a protofilament and a GCP-spoke of γ-TuRC-capped microtubule minus-end. (**A**) Cross-sectional view showing “luminal-bridge” density (orange). Stable (solid line) and dynamic (dashed line) regions are indicated. (**B, C**) Example of the γ-tubulin density (mesh) from the stable region, along with a schematic of γ-TuRC indicating the most stable positions in blue. Rigid body fitted model (sticks) for H12 helix (B) and GTPase domain β-sheets (C). (**D**) An ɑ/β-tubulin dimer bound to γ-tubulin, positioned above a GCP (backbone: ribbons, nucleotide: spheres). (**E**) Inset from (D) shows an overlay of cryo-EM density (mesh) with guanine nucleotide coordinates (stick representation) found at γ/ɑ-tubulin interface. (**F-H**) Structural regions of γ-tubulin, ɑ-tubulin and GCP (spheres) interacting with ɑ-tubulin, β-tubulin and γ-tubulin (surface), respectively. (**I**) Overlaying protofilament density at position 7 (transparent gray surface) with rigid-body fitted molecular model reveals structural alignment extending from GCP, through γ-tubulin, to an ɑ/β-tubulin dimer.

Refinements using signal subtraction and focused classification without any averaging or applied symmetry for the stable region yielded a ∼7.5Å resolution map (materials and methods; fig S2), sufficient for rigid-body fitting of individual domains and enabling detailed inspections of key protein-protein interfaces (e.g. γ-tubulin-α-tubulin). As significant differences were not observed across all positions, we then applied symmetry expansion and averaged the densities of the most stable positions (positions 5-9) to generate an improved focused map of two γ-tubulins bound to two α-tubulins (resolution ∼4.0Å) (materials and methods; fig S2), sufficient to visualize twists within single helices (Fig. 2B) and individual strands of β-sheets (Fig. 2C), for globular densities and were able to rigid body fit models and assign γ- and α-tubulins. For position 7, we isolated a single spoke of γ-TuRC and the associated protofilament and built a model for GCP2, γ-tubulin and α/β-tubulins (Fig. 2D).

These models reveal that the γ- and α-tubulin interface contains the nucleotide binding site in γ-tubulin (Fig. 2D). At the current resolution, we were able to identify that the pocket is occupied with a nucleotide (Fig. 2E), however, we are unable to distinguish between GDP vs GTP. This interface also involves three additional contacts spanning the width of γ- and α-tubulin (Fig. 2F). Notably, these contacts are reminiscent of the main contacts that establish the α- and β-tubulin interfaces (Fig. 2G). Remarkably, we also observed three similar contacts between the GCP-C-terminal GRIP2 domain and γ-tubulin interface (Fig. 2H), revealing structural mimicry.

In addition, we observed structural alignment, extending from γ-TuRC to the α/β-tubulins in the protofilaments, involving two secondary structure features. First, helices (H3-H4) of the α-, β- and γ-tubulins align vertically with helices in the GCP ‘notch’ (Fig. 2I). Second, the C-terminal domains of α-, β- and γ-tubulins align with GCP-C-terminal helices. The spacing between successive α-tubulins within a protofilament is ∼82Å, consistent with a compacted lattice form (*32*). Importantly, the spacing between γ-tubulin and the first β-tubulin vertically above is also ∼82Å. Together, our findings reveal unanticipated structural mimicry and longitudinal alignments of key secondary structure elements between α/β-tubulins and the microtubule-bound γ-TuRC.

### The γ-TuRC undergoes extensive structural rearrangements at the overlap region to match the geometry of the microtubule lattice

To examine how γ-TuRC, which has 14 γ-tubulins binds a microtubule with 13 protofilaments, we focused on the dynamic region of γ-TuRC. We performed additional classifications focusing on positions 1 and 11-14 of the microtubule-bound γ-TuRC (materials and methods; fig S2). We identified a subset of particles (8,195 ends) that had stronger densities for the GCP spokes at the dynamic region (Fig. 2A). Further processing and refinement using this subset of particles yielded a map (overall resolution of 9.5Å) that revealed stronger densities at γ-TuRC’s overlap region (Fig. 3A,B). This reconstruction revealed how a globular density, consistent with the actin in the luminal bridge (orange), acts as a steric constraint to help position the GCPs (pink) that overlap in the γ-TuRC (Fig. 3B).

**Fig. 3.**
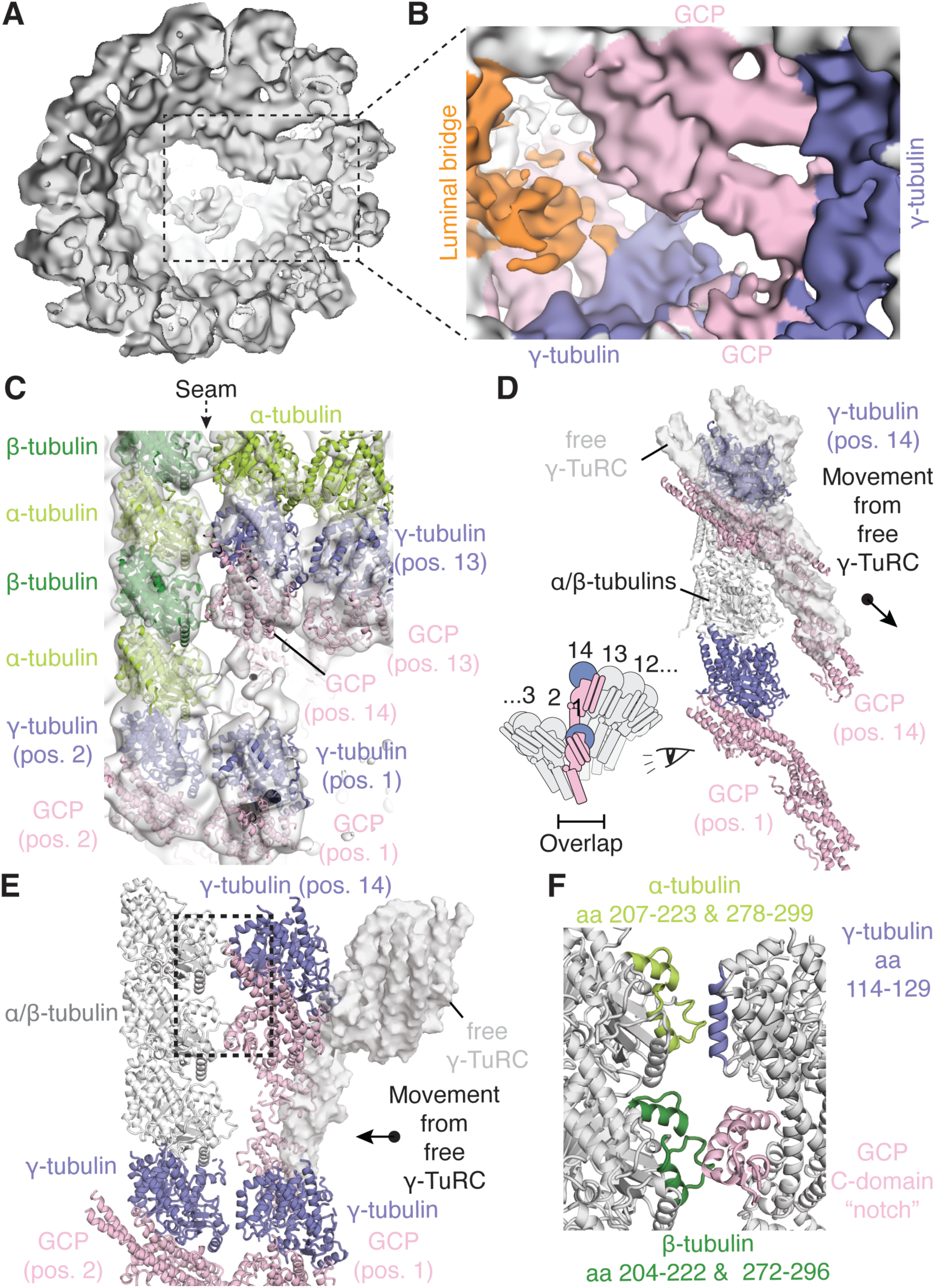
Structural rearrangements of tubulins and GCPs at the seam. (**A**) Surface representation of overall reconstruction (lowpass-filtered to 13 Å) for γ-TuRC-capped microtubule using a subset of particles with stronger densities at the dynamic region. (**B**) Zoomed-in view of the dynamic region (viewed from the luminal side, colored as indicated). (**C**) Rigid-body fit of ɑ,β- and γ-tubulins and GCP domains into the density (lowpass-filtered to 13 Å, transparent gray surface) at the dynamic region. (**D-E**) Side- and front views of the seam showing γ-TuRC-capped microtubule models (ribbons) and free γ-TuRC model (transparent gray surface). (**F**) Inset from (E) highlighting the interacting structural motifs at the seam for the α-tubulin-γ-tubulin interface and the GCP-β-tubulin interface.

We next determined a structure of recombinant γ-TuRC using the microtubule-free complexes in our micrographs (Fig. 1E and fig. S2). Cleaned free γ-TuRC particles were processed and classified to select a subset of ∼150,000 particles with the strongest signal for the overlap region. These particles were refined to generate a density map at an overall resolution of ∼7.3Å and sufficient for rigid-body docking of individual domains. The overall γ-TuRC structure was similar to that of the native complex (*8*) (fig. S4A,B). Rigid-body fitting of the free γ-TuRC molecular model into γ-TuRC-bound microtubule structure revealed extensive mismatch, suggesting that there were substantial rearrangements across the entire microtubule-bound complex. The greatest mismatch was apparent at the overlap region.

To account for this structural mismatch we rigid-body docked individual segments, composed of GCP, γ-tubulin, and α/β-tubulins from position 7, into all positions in the density map for the microtubule-bound γ-TuRC revealing significant rearrangement of the complex (materials and methods; fig. S4C,D). A view of this model from the exterior revealed that γ-tubulin from position 14 is ∼12 nm above the γ-tubulin from position 1 of the γ-TuRC (Fig. 3C). Remarkably, both these γ-tubulins are aligned vertically to each other and to the first protofilament in the microtubule (above position 2). Below each of these γ-tubulins, we observe densities for the GCP-C-terminal domains and a gap (∼4 nm) between the GCP-C-terminal domain from position 14 and the γ-tubulin from position 1 in γ-TuRC. In addition, as α-tubulin and γ-tubulin interact head-to-tail (Fig. 1I and Fig. 2D,F), the α-tubulin above the position 14 γ-tubulin laterally interacts with the β-tubulin above γ-TuRC’s position 2 to establish the seam of a 13-protofilament microtubule with α/β-tubulins arranged in a 3-start helix (Fig. 3C).

To examine this arrangement of the γ-tubulins in positions 1 and 14, we compared the free- and microtubule-bound γ-TuRC models. Aligning spokes in positions 1 of the two γ-TuRC models revealed that upon microtubule binding the GCP C-domain in position 14 is ‘tucked’ further into the lumen (Fig. 3D) and above the GCP in position 1. The γ-tubulin and GCP-C-terminal domain in positions 14 is also displaced towards position 2, so that it is proximal to the microtubule lattice (Fig. 3E). In particular, the γ-tubulin from the 14th position is in lateral contact with α-tubulin, which is above an α/β-tubulin dimer directly above position 2 of the γ-TuRC. This interface between γ-tubulin and α-tubulin, which has not been previously observed (Fig. 3F), mimics the interactions between lateral tubulin-tubulin interactions within the microtubule lattice. In addition, lateral contacts are also formed between the GCP-C-terminal GRIP2 domain and β-tubulin, predicted by an extrapolated model based on γ-TuSC assemblies (*17*) (Fig. 3F). Specifically, these interactions involve a ‘notch’ formed from a GCP GRIP2 domain and β-tubulin (aa:204-222 and aa:272-296). Together, these reconstructions reveal the extensive rearrangements of the γ-TuRC subunits that are stabilized by new protein-protein interfaces to align the overlap region of the γ-TuRC to the seam of a 13-protofilament microtubule.

### The entire γ-TuRC compacts to match the 13-protofilament microtubule lattice

Our reconstruction of the stable portion of the γ-TuRC-capped microtubule (positions 3-9), followed by symmetry expansion reveals how γ-tubulins in the microtubule-bound γ-TuRC interact laterally (materials and methods; Fig. 4A and fig. S2). We find that this interface is established by two structural features. First, the M-loop (S7-H9 loop) in a γ-tubulin interacts with a cleft formed by H1-S2 and H2-S3 loops (Fig. 4B). Second, an additional interface is formed between the helix H3 and H3-S5 loop from one γ-tubulin and helices H9-H10 in the adjacent γ-tubulin (Fig. 4C). These two contacts are structurally equivalent to those between α-tubulin-α-tubulin and β-tubulin- β-tubulin in the microtubule lattice (fig. S3).

**Fig. 4.**
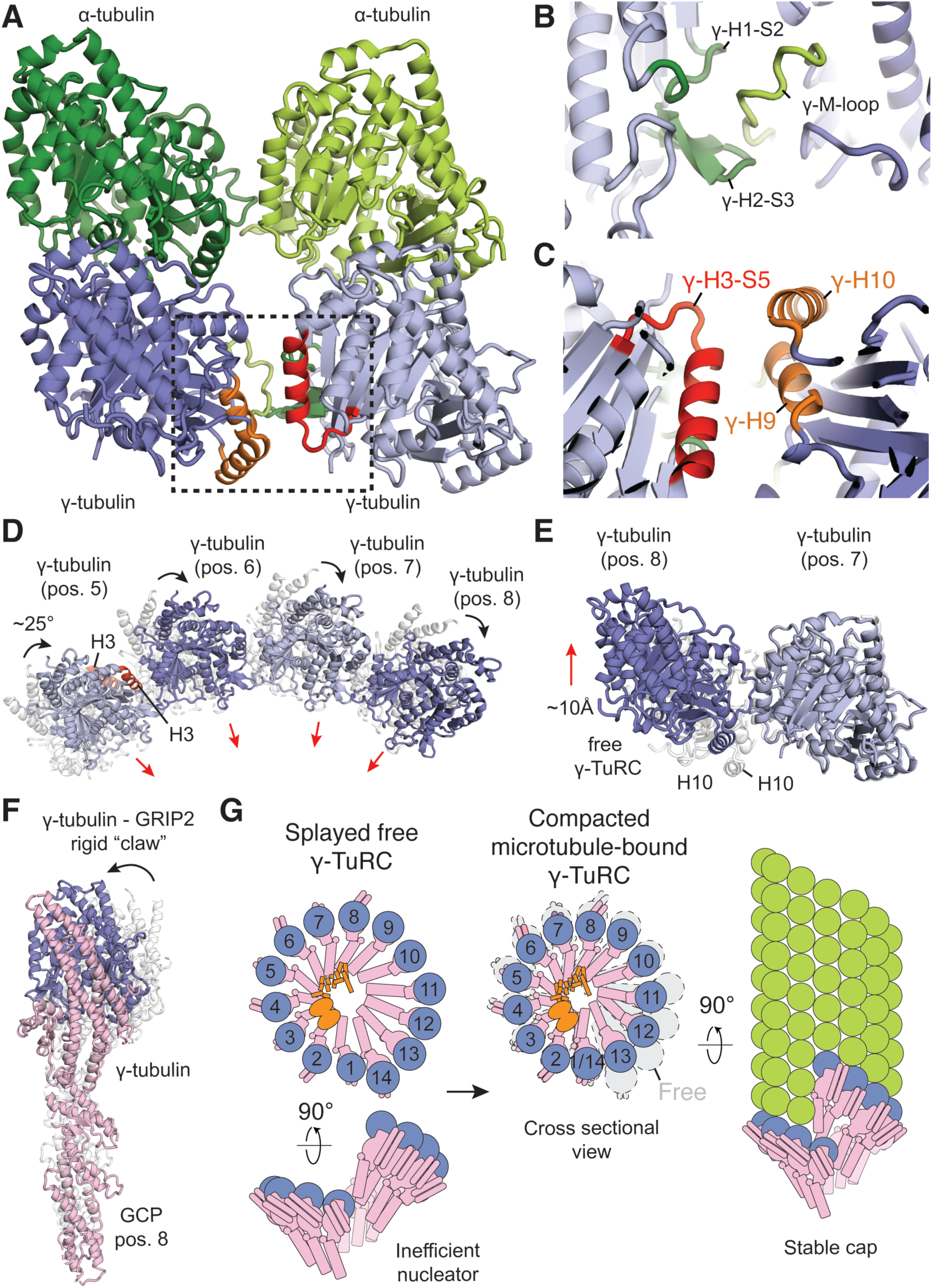
Lateral interactions established upon the compaction of γ-TuRC to cap the microtubule minus-end. (**A**) Lateral interactions between ɑ-ɑ- and γ-γ-tubulins (ribbons), with two key interaction motifs between adjacent γ-tubulins highlighted (shades of green and red). (**B,C**) Inset from (A) shows these two structural motifs in detail. (**D**) Overlay of γ-tubulin in the free (ribbons: gray) and microtubule-bound (ribbons: shades of blue) γ-TuRC. Arrows highlight rotation and displacements of the γ-tubulin’s. The H3 helix (light red: free; dark red: microtubule-bound) is indicated. (**E**) Vertical displacements of the γ-tubulins in the microtubule-bound complex (shades of blue) relative to the free complex (gray). The H10 helix in γ-tubulin is indicated. (**F**) The rigid-body motion between the free (gray) and microtubule-bound (colored) states of γ-tubulin–GRIP2-domain of GCP, which together form a rigid “claw”, is indicated. (**G**) Schematic for how γ-TuRC, an inefficient nucleator (splayed complex) transforms into a stable microtubule minus-end cap (compact complex), GCP spokes are colored in pink, γ-tubulins in purple, the luminal-bridge components in orange and ɑ/β-tubulins in green.

This microtubule lattice-compatible γ-tubulin-γ-tubulin interface is not observed in free γ-TuRC and the helical γ-TuSC structures (*8*, *9*, *23*). In free γ-TuRCs, each γ-tubulin can establish longitudinal contacts with α-tubulin (Fig. 2). However, in free γ-TuRCs the γ-tubulins have two alternating spacings (9 and 15Å) and are rotated such that the lateral contacts we observe in the microtubule-bound complex are not possible (*8*). For example, the H3 helix is pointing away from the center of the helical axis, rather than being oriented to contact the adjacent γ-tubulin (Fig. 4D). Overlays of the γ-tubulins in the free and microtubule-bound γ-TuRC reveal that each of the γ-tubulins undergo ∼25 degree rotation and a displacement towards the middle of the complex, resulting in overall compaction of the complex (Fig. 4D and fig. S4C,D). Further, the vertical displacement of the γ-tubulins at the interface relative to each other is increased (∼10Å) in the microtubule-bound complex compared to the free complex (Fig. 4E). These vertical and lateral displacements result in highly uniform positioning of γ-tubulins in the microtubule-bound γ-TuRC (Fig. 4E).

Comparisons of the free and microtubule-bound structures indicate that there are no substantial movements across the GCP N-domains (positions 1-12), which mediate interactions between the GCPs and coalesce to form the base of the cone-shaped complex (Fig. 1G-I). Remarkably, the lateral contacts are altered such that the GRIP2 domain, which interacts with the adjacent γ-tubulin in the free complex, is displaced to allow lateral contacts between adjacent γ-tubulins in the microtubule-bound complex (Fig. 4A). Strikingly, γ-tubulin and the GCP-GRIP2 domain form a rigid ‘claw’ that moves towards the γ-TuRC lumen and rotates in the direction of the helical rise (Fig. 4F) (*17*). Together, our findings reveal how the γ-TuRC compacts to establish extensive new interactions needed to match the 3-start helical geometry of a 13-protofilament microtubule.

## DISCUSSION

We show that the γ-TuRC selectively nucleates 13-protofilament microtubules. Our data also provide direct structural evidence for the ‘template’ model of γ-TuRC-dependent nucleation and reveal how the even number of γ-tubulins in γ-TuRC align with the odd number of α-tubulins at the microtubule minus-end. Compared to non-13-protofilament microtubules, which provide intracellular transport tracks with super helical twists, 13-protofilament microtubules provide tracks aligned to the filament’s long axis. This lattice feature is likely important not only for directional motor protein-dependent transport, but also for microtubule crosslinking and relative sliding. Helical twists in the trajectories of crosslinking motor proteins could result in increased mechanical strain and constraints during the sorting of two or more microtubules into bundles (e.g. parallel kinetochore fibers, A-tubules in motile cilia and anti-parallel midzone bundles) that have essential force-generating cellular functions (*33–35*).

Our data show how nanometer-scale compaction of the γ-TuRC is achieved through motions in each of its 14 spokes, revealing how these small but additive inter- and intra-domain movements can regulate microtubule lattice architecture on the micrometer scale (Fig. 4G). The longitudinal contacts between α- and β-tubulins along a protofilament have been found to be stronger than the lateral contacts between protofilaments (*36*, *37*), suggesting that γ-TuRC would position γ-tubulins to promote and stabilize these lateral interactions. This γ-tubulin positioning requires all the GCPs in γ-TuRC reorient, with the GRIP2 domain and γ-tubulin forming a rigid ‘claw’ that pivots relative to the stable GCP N-terminal domains. This large-scale compaction of the complex accommodates the actin-containing luminal bridge, which stabilizes and closes the ring of the free complex (*24*, *38*). The extensive conformational changes needed to transform γ-TuRC from the free state to the microtubule-bound state are likely to be why this complex is an inefficient microtubule nucleator (Fig. 4G). Importantly, these structural data suggest how nucleation may be regulated through promoting these structural changes or stabilizing conformations that more closely match or pre-organize γ-TuRC.

Our findings explain why the γ-TuRC functions as an effective minus-end cap that can remain bound to the newly formed microtubule minus end for several minutes (*25*). At least three factors likely contribute to this stable end-capping. First, in the microtubule-bound γ-TuRC conformation, the lateral interactions between adjacent γ-tubulins mimic those between α- and β-tubulins pairs and do not result in substantial deformations of the microtubule minus-end. Second, the overall structural similarity between γ-tubulin and α-tubulin allows for these two tubulins to interact at the overlap region of the γ-TuRC to establish the ‘seam’ of the microtubule. Third, the lateral interactions between GCP’s GRIP2 domain with β-tubulin at the seam further stabilize this interface. Importantly, the longitudinal interactions in the microtubule lattice, which are modulated by nucleotide states or drug binding (*36*, *37*), remain compatible with the γ-TuRC-binding to the minus end.

Together, our findings provide a mechanistic framework to explain the biochemical and modeling data collected over decades for how 13-protofilament microtubules are templated by the γ-TuRC and how microtubule minus-ends are capped. Our on-grid nucleation approach should allow structural analyses to extend beyond that of the microtubule lattice and interacting proteins (e.g. kinesins) to more complex processes (e.g. branching or anchoring at minus-ends) that are essential for network dynamics and function.

## METHODS

### Molecular Cloning

Recombinant γ-TuRC plasmids used were identical to the plasmids used in a previous study (*24*). To express recombinant tubulin, we modified the insect cell codon-optimized expression constructs for human tubulin comprising of the ɑ-β tubulin isoforms: ɑ-tubulin-TUBA1B (NP_006073.2) and β-tubulin TUBB3 (NP_006077.2) from (*39*) using gibson assembly based on a strategy described in (*31*). Briefly, the sequence encoding the deca-histidine tag, Tobacco Etch Virus (TEV) protease site and the alanine-proline linker at the N-terminus of ɑ-tubulin were removed. A hexa-histidine tag encoding DNA sequence was inserted between isoleucine 42 and glycine 43 in the acetylation loop. The resulting expression construct was: TUBA1B-internal hexa-histidine tag TUBB3-Gly-Gly-Ser-Gly-Gly linker-TEV site-Strep-tag II, referred to as wild type tubulin in the manuscript. This construct was further subjected to mutagenesis using gibson assembly to generate the previously reported GTP-hydrolysis compromised TUBA1B (E254D)-internal hexa-histidine-tag TUBB3-Gly-Gly-Ser-Gly-Gly linker-TEV site-Strep-tag II construct (*31*), referred to as E254D tubulin in the manuscript.

chTOG construct used was the same as described in (*40*).

### Purification of native γ-TuRC

Native γ-TuRC was purified with a slight modification of the published protocol from (*8*). Briefly, native γ-TuRC was purified from HeLa S3 cytoplasmic extracts using the affinity of human γ-TuRC for the CM1 region or γ-TuRC-mediated nucleation activator (γ-TuNA) motif of CDK5RAP2, which has been reported to act as a γ-TuRC-activator (*41*), followed by sucrose density gradient centrifugation. The HIS_6_-SUMO-TEV-GFP-PreScission-γ-TuNA expression construct in the modified pET plasmid from (*8*), encoding the amino acid residues 51-100 corresponding to the CM1 region of human CDK5RAP2, was transformed into BL21(DE3) Rosetta cells (Novagen). γ-TuNA expression was induced in a 12 L culture with 0.5 mM IPTG for 16 hours at 18°C. The bacteria were harvested by centrifugation at 5,000 g for 10 minutes at 4°C and resuspended in 240 mL lysis buffer (50 mM sodium phosphate, 300 mM NaCl, 15 mM Imidazole, 0.1% (v/v) Tween-20, 1 mM 2-Mercaptoethanol and 6 cOmplete EDTA-free protease inhibitor Cocktail tablets (Roche), pH 8.0). The cells were lysed by passing them through an Emulsiflix C-5 (Avestin) homogenizer three times followed by clarification at 35,000 rpm, for 1 hour at 4°C in a Type 45 Ti rotor (Beckman Coulter). The clarified lysate was then incubated with 5 mL of Ni-NTA agarose beads (Qiagen), pre-equilibrated in lysis buffer, for 1 hour at 4°C in the cold room. Following extensive washing in lysis buffer, the protein was eluted in Ni-NTA elution buffer (50 mM sodium phosphate, 300 mM NaCl, 300 mM Imidazole, 0.1% (volume/volume) Tween-20, 1 mM 2-Mercaptoethanol, pH 8.0). Protein-containing fractions were identified by Bradford assay, pooled together and concentrated down to ∼2 mL with an Amicon Ultra 10K MWCO centrifugal filter unit (Millipore UFC901024). The HIS_6_-SUMO-tag was cleaved by proteolysis with 1 mg TEV protease for 2 hours at 4°C in the cold room. The protein was further purified by size-exclusion chromatography through a HiLoad 16/60 column Superdex 75 column, pre-equilibrated in gel filtration buffer (40 mM HEPES, 150 mM NaCl, 1 mM MgCl_2_, 2 mM 2-Mercaptoethanol, pH 7.5) . Peak fractions containing γ-TuNA-GFP were identified by SDS-PAGE, pooled together and the protein was snap-frozen in liquid nitrogen after supplementing it with 10% (weight/volume) sucrose and stored at -80°C.

GFP-nanobody, used for making the nanobody column, was purified in the same way as reported (*8*, *42*). Briefly, periplasmic expression of the recombinant GFP-nanobody in BL21(DE3) Rosetta cells (Novagen) was induced with 0.1 mM IPTG for 16 hours at 16°C . Cells from a 6 L culture of the bacteria were harvested by centrifugation at 5,000 g for 10 minutes at 4°C and resuspended in 60 mL ice-cold TES buffer (200 mM Tris-Cl, 0.5 mM EDTA, 500 mM Sucrose, pH 8.0). After resuspension using vortexing, the cells were osmotically shocked on ice for 30 minutes by diluting them 5-fold in a 1:4 (volume/volume) mixture of ice-cold TES buffer:water. Periplasmic fraction was then isolated first by a low speed centrifugation at 6,000 g for 10 minutes to get rid of cell debris and then a second spin at 20,000 g for 20 minutes at 4°C. The supernatant from the second spin was incubated with 3 mL of Ni-NTA agarose beads (Qiagen) for 30 minutes at 4°C in the cold room. The beads were subsequently washed with wash buffer A (20 mM Tris-Cl, 900 mM NaCl, pH 8.0), followed by a wash with wash buffer B (20 mM Tris-Cl, 150 mM NaCl, 10 mM Imidazole, pH 8.0). The nanobody was then eluted with Ni-NTA elution buffer (20 mM Tris-Cl, 150 mM NaCl, 250 mM Imidazole, pH 8.0), peak fractions were identified by Bradford assay, pooled together and concentrated down to 0.5 mL with a Amicon Ultra 10K MWCO centrifugal filter unit (Millipore UFC901024). Finally, the GFP nanobody was purified by gel filtration over a Superdex 75 10/300 column, pre-equilibrated with gel filtration buffer (100 mM sodium bicarbonate, 150 mM NaCl, pH 8.0). Peak fractions were pooled together and ∼10 mgs of GFP nanobody was conjugated to a 1 mL NHSTrap column (Cytiva) according to manufacturer’s protocol.

Approximately 45 mL of frozen cytoplasmic extract from 10 L of HeLa S3 cells (*43*), a kind gift from Dr. Robert Roeder, was first rapidly thawed partially in room temperature water and subsequently completely thawed on ice. The extract was diluted 1:1 with 45 mL of ice-cold γ-TuRC buffer (50 mM HEPES, 150 mM KCl, 1 mM MgCl_2_, 1 mM EGTA, 0.1% (volume/volume) IGEPAL and 0.1 mM GTP, 1 mM PMSF, 3 U/mL of Benzonase, 1 cOmplete EDTA-free protease inhibitor Cocktail tablet (Roche), 1 mM DTT, pH 7.5). The diluted extract was then centrifuged at 31,200 rpm for 30 minutes at 4°C in a Type 70 Ti rotor (Beckman Coulter). The supernatant was filtered through a 0.45 µm syringe filter (Millex-GP PES membrane (Millipore SLHP033RS) and incubated with 15 mgs of GFP-γ-TuNA for 1 hour on ice in the cold room. The extract and γ-TuNA mixture was then loaded onto a GFP nanobody column, followed by washes with γ-TuRC buffer, without protease inhibitors until the UV absorbance reached baseline. The GFP tag was then cleaved off with PreScission protease (∼3 mgs), diluted in γ-TuRC buffer and injected onto the nanobody column. The proteolysis was allowed to proceed for 6 hours on the column, following which γ-TuRC was eluted with γ-TuRC buffer. Peak fractions were pooled together and loaded onto a 2 mL sucrose gradient composed of 5%, 16.7%, 28.3% and 40% (weight/volume) sucrose in gradient buffer (40 mM HEPES, 150 mM KCl,1 mM MgCl_2_, 1 mM EGTA, 0.01% (volume/volume) IGEPAL, 0.1 mM ATP, 0.1 mM GTP, 1 mM DTT, pH 7.5) in a 11 x 34 mm Ultra-Clear centrifuge tubes (Beckman Coulter: 347356). The sucrose density gradient was centrifuged at 50,000 rpm for 3 hours at 4°C in a TLS-55 swinging bucket rotor (Beckman Coulter) with minimum acceleration up to 5,000 RPM and deceleration with no break. The gradient was manually fractionated in the cold room using a P-1000 cut tip into 150 µL fractions and the fractions were analyzed by negative stain electron microscopy for the presence of lock-washer shaped complexes. Peak fractions, based on a higher number of γ-TuRC complex particles per micrograph, were pooled together and snap-frozen in liquid nitrogen and stored at -80°C.

### Expression and purification of recombinant tubulin

Donor plasmids encoding the wild type or mutant tubulin were transformed into Multi-bac Turbo cells (ATG:biosynthetics GmbH) and positive colonies were selected based on blue/white screening. Isolated bacmids were transfected into Sf9 cells (Novagen) using the Bac-to-Bac system (Life Technologies) to generate the baculovirus. Baculoviruses were amplified twice and freshly generated P3 virus was used to infect 1.8-2 L of High Five cells (Thermo Fisher Scientific) at a density of 3-3.5 x10^6^ cells/mL by adding 6 mL virus per 600 mL of cells. Cells were harvested after 60 hours post infection at 27°C by centrifugation at 1,000 g for 10 minutes at 4°C. Cells were lysed in 60 mL of ice-cold lysis buffer (50 mM HEPES, 20 mM imidazole, 100 mM KCl, 1 mM MgCl_2_, 0.5 mM β-mercaptoethanol, 0.1 mM GTP, 3 U/mL benzonase, 3 cOmplete EDTA-free protease inhibitor Cocktail tablets (Roche), pH 7.2) by dounce homogenization (20 strokes).The lysate was clarified by centrifugation at 56,000 rpm in a Type 70 Ti rotor (Beckman Coulter) at 4°C for 1 hour, filtered through 0.22 µm syringe filter (Millex-GP PES membrane (Millipore SLGP033RS) and then loaded onto a 5 mL HisTrap HP column (Cytiva), previously equilibrated in lysis buffer. The column was then washed with 40 mL of lysis buffer or until the UV absorption reached baseline and bound protein was eluted with nickel elution buffer (1X BRB80 (80 mM PIPES, 1 mM MgCl_2_, 1 mM EGTA), 500 mM imidazole, 0.2 mM GTP, 2 mM β-mercaptoethanol, pH 7.2). Protein-containing fractions were pooled together (∼15 mL), diluted 3-fold with lysis buffer and loaded onto two 1 mL StrepTrap XT columns (Cytiva), connected in tandem, using a Superloop. The columns were washed with 25 mL of 66% lysis buffer + 34% nickel elution buffer mixture, followed by 25 mL of wash buffer A (1X BRB80, 1 mM β-mercaptoethanol, 0.1 mM GTP, 0.1% (volume/volume) Tween-20, 10% (weight/volume) glycerol, pH 7.2) and finally 25 mL of wash buffer B (1X BRB80 1 mM β-mercaptoethanol, 0.1 mM GTP, 10 mM MgCl_2_, 5 mM ATP, pH 7.2). The column-bound protein was eluted with StrepTrap XT elution buffer (1X BRB80, 20 mM Imidazole, 2 mM β-mercaptoethanol, 0.2 mM GTP, 50 mM biotin, pH 7.2). Protein fractions eluted from StrepTrap XT elutions were pooled together and mixed with 4 mg of TEV protease (∼7 mg/mL stored in 40 mM HEPES, 150 mM KCl, 30% (weight/volume) glycerol, 1 mM MgCl_2_, 3 mM β-mercaptoethanol, pH 7.5, diluted into 5 ml StrepTrap XT elution buffer). Proteolysis was carried out for 2 hours on ice. Post-TEV digestion, the protein was loaded onto two 1 mL HiTrap SP columns (Cytiva), followed by a wash with StrepTrap XT elution buffer. The flow-through of the SP columns, containing tubulin was pooled, concentrated with Amicon Ultra 50K MWCO centrifugal filter unit (Millipore UFC901024), and loaded on to a Superdex 200 16/60 column (GE life science 17-1069-01), previously equilibrated in size-exclusion buffer (1X BRB80, 5% (weight/volume) glycerol, 0.2 mM GTP, 2 mM β-mercaptoethanol, pH 6.8). Peak fractions containing tubulin were identified by SDS-PAGE, pooled together and concentrated to ∼5 mg/mL with an Amicon Ultra 50K MWCO centrifugal filter unit. The concentrated protein was snap-frozen in liquid nitrogen and stored at -80°C.

### Expression and purification of recombinant γ-TuRC

Recombinant γ-TuRC holo-complex was purified based on a previously published protocol (*24*) using ZZ-TEV-MZT2-mEGFP as an affinity handle for the complex. Briefly, donor plasmids encoding for all 10 components of γ-TuRC (pAceBac1-γ-TuRC-GFP) or just γ-tubulin-TEV-HIS, GCP2 and GCP3 (pAceBac1-γ-TuSC) were transformed into Multi-BacTurbo cells (ATG:biosynthetics GmbH). Positive colonies, based on blue/white screening were used to generate bacmids, which were transfected into Sf9 cells (Novagen) using the Bac-to-Bac system (Life Technologies) to generate the corresponding γ-TuRC-GFP and γ-TuSC baculoviruses. Following two rounds of baculovirus amplification, fresh P3 virus from γ-TuRC-GFP and γ-TuSC were mixed in a 1:1 ratio and used to infect 1.8-2 L of High five cells (Thermo Fisher Scientific) at a density of 3-3.5 x10^6^ cells/mL by adding 30 mL of the γ-TuRC-GFP and γ-TuSC virus mixture as above, per 600 mL of cells. 60 hours post infection at 27°C, High five cells were harvested by centrifugation at 1,000 g for 10 minutes at 4°C. The cell pellet was resuspended in 80 mL of lysis buffer (40 mM HEPES, 150 mM KCl,1 mM MgCl_2_,10% [volume/volume] glycerol, 0.1% [volume/volume] Tween-20, 0.1 mM ATP, 0.1 mM GTP, 1 mM 2-mercaptoethanol, pH 7.5, four cOmplete EDTA-free Protease Inhibitor Cocktail tablets (Roche), 500 U benzonase, 2 mM PMSF, and 4 mM benzamidine-HCl) and lysed by dounce homogenization on ice (20 strokes). The lysate was clarified by centrifugation at 56,000 rpm in a Type 70 Ti rotor (Beckman Coulter) at 4°C for 1 hour. Clarified lysate was filtered through 0.22 µm syringe filter (Millipore SLGP033RS) and loaded onto a 1 mL NHSTrap column (Cytiva), which was previously coupled to 10-20 mg rabbit IgG (Innovative Biosciences IR-BIGGAP500 MG) as per manufacturer’s protocol. The column was washed with 30 mL of lysis buffer followed by 10 mL of gel filtration buffer (40 mM HEPES, 150 mM KCl, 1 mM MgCl_2_, 10% (volume/volume) glycerol, 0.1 mM GTP and 1 mM 2-mercaptoethanol, pH 7.5,). On-column proteolysis was done with 1 mg TEV protease (∼7 mg/ml stored in 40 mM HEPES, 150 mM KCl, 30% (weight/volume) glycerol, 1 mM MgCl_2_, 3 mM β-mercaptoethanol, pH 7.5), diluted into 1 mL gel filtration buffer for 2 hours at 4° C. The eluate containing the digested protein was pooled together and concentrated by dialysis with the dialysis buffer (40 mM HEPES,150 mM KCl,1 mM MgCl_2_, 60% [weight/volume] sucrose, 0.1 mM GTP and 2 mM 2-mercaptoethanol, pH 7.5) for 4 hours at 4°C. The concentrated protein was loaded onto a Superose 6 Increase 10/300 GL column (Cytiva), previously equilibrated in gel filtration buffer. Peak fractions from the first peak, corresponding to γ-TuRC-GFP were pooled together and loaded onto a 2 mL sucrose gradient composed of 10%, 20%, 30% and 40% sucrose in gradient buffer (40 mM HEPES, 150 mM KCl,1 mM MgCl_2_, 0.01% (volume/volume) Tween-20, 0.1 mM ATP, 0.1 mM GTP, 1 mM 2-mercaptoethanol, pH 7.5). Sucrose density gradient centrifugation was done by centrifugation at 50,000 rpm for 3 hours at 4° C. in a TLS-55 swinging bucket rotor (Beckman Coulter) with minimum acceleration upto 5,000 rpm and deceleration with no break. The gradient runs were manually fractionated using a P-1000 cut tip into 250 uL fractions and analyzed by SDS-PAGE followed by Coomassie staining and negative stain electron microscopy. Peak fractions corresponding to the highest amount of protein by Coomassie were pooled together, aliquoted and snap-frozen in liquid nitrogen. Protein aliquots were stored in -80 until further use.

### Negative stain-EM

3.3 µl of sucrose density gradient–purified native γ-TuRC fraction was applied to carbon-coated copper grids (EMS; CF-400-Cu) glow-discharged for 1 minute. The protein solution was incubated on the grid for 1 minute at room temperature, then manually blotted away by gently touching the rim of the grid with a Whatman 1 filter paper (Whatman, CAT No.1001-090) to wick away the unbound protein solution and replaced with another 3.5 µL. This was repeated for a total of three applications. Post the final application, the protein solution was manually blotted away by gently touching the rim of the grid on one side with a Whatman 1 filter paper and freshly filtered 1% (weight/volume) uranyl acetate was simultaneously applied from the opposite side to exchange the solution. The grid was incubated in uranyl acetate for a 1 minute. The stain was removed by manual blotting, and grids were air-dried. Micrographs were collected on an FEI Tecnai G2 microscope operating at 120 kV at a magnification of 30,000× (3.036 Å/pixel) using a BioSprint 29 charge-coupled device camera.

### Cryo-EM sample preparation

#### Native γ-TuRC nucleated microtubules sample

To get a higher number of native γ-TuRCs per micrograph, we used the multiple application technique (*44*). 1.2 µL of native γ-TuRC aliquot from the sucrose gradient fraction was applied to a Lacey Carbon Support Film grid with an Ultrathin Carbon Film (< 3 nm) (TED PELLA, INC, Prod No. 01824) or Quantifoil R2/2 300-square-mesh copper grids coated with continuous carbon film, plasma cleaned for 30 s. After 3 minutes of incubation, the sample was manually blotted away by gently touching the rim of the grid with a Whatman 1 filter paper (Whatman, CAT No.1001-090) to wick away the unbound protein solution and replaced with another 1.2 µL of protein. This was repeated for 5 applications in total. 10 µL of assay buffer (1X BRB80 [80 mM PIPES, 1mM MgCl_2_, 1mM EGTA], 1 mM GTP, 50 mM KCL, 1 mM DTT, pH 6.8, adjusted with KOH, 0.15% (weight/volume) methyl-cellulose and 0.2 mg/mL κ-casein, prepared in 1X BRB80), was then applied onto the grid to get rid of the sucrose in the γ-TuRC gradient buffer. Immediately after applying the assay buffer, the tweezer and the grid were moved to a closed chamber on a heat block at 37°C for 5 minutes. The assay buffer was wicked away with a Whatman 1 filter paper and replaced with 3.3 µL of nucleation/polymerization mixture containing 20 µM of cycled bovine brain tubulin, 100 nM chTOG and and oxygen scavenging system (0.35 mg/mL catalase, 0.2 mg/mL glucose oxidase, 2.5 mM glucose, and 10 mM DTT) prepared in assay buffer. After 2 minutes of incubation at 37°C, the nucleation/polymerization mixture was replaced with another 3.3 µL of the same. This was repeated 3 times for a total incubation time of 6-9 minutes with the nucleation/polymerization mixture. Next, in order to get rid of methyl cellulose, the grid was incubated with a nucleation/polymerization mixture without methyl-cellulose for 1 minute at room temperature. The grid was then transferred to the vitrobot tweezer and then to the Vitrobot IV (Thermo Scientific) arm, pre-equilibrated to 37°C. The grid was blotted for 3-3.5 s at 100% humidity and 37°C and then plunge frozen in liquid ethane.

Data was collected on 5 grids on FEI Titan Krios microscopes using Serial EM automated data collection (*45*). The data acquisition parameters are summarized in Table S1.

#### Spontaneously nucleated microtubules sample

For microtubules nucleated spontaneously in solution, the sample for cryo-EM was prepared in a similar way as above without the step for native γ-TuRC application on the grid. 5 uL of nucleation/polymerization mixture containing 20 µM of cycled bovine brain tubulin, 100 nM chTOG and and oxygen scavenging system (0.35 mg/mL catalase, 0.2 mg/mL glucose oxidase, 2.5 mM glucose, and 10 mM DTT) prepared in assay buffer, was applied on to Quantifoil R2/2 300-square-mesh copper grids coated with continuous carbon film, freshly plasma cleaned for 30 s. The grid was incubated in a closed chamber on a heat block at 37°C for 45 minutes. The grid was removed from the closed chamber and incubated with a nucleation/polymerization mixture without methyl-cellulose for 1 minute at room temperature. The grid was subsequently transferred to vitrobot tweezer and then to the Vitrobot IV (Thermo Scientific) arm, pre-equilibrated to 37°C. It was blotted for 3s at 100% humidity and 37°C and then plunge frozen in liquid ethane.

Data was collected in a session on a FEI Titan Krios microscope using Serial EM automated data collection (*45*).

#### Recombinant γ-TuRC nucleated microtubules sample

In order to have γ-TuRC particles at a high density on the grid, multiple application technique (*44*) was used. 3.5 µL of a γ-TuRC aliquot from sucrose gradient was applied to Lacey Carbon Support Film grid with an Ultrathin Carbon Film (< 3 nm) (TED PELLA, INC, Prod No. 01824), plasma cleaned for 30 s, held in place with the vitrobot tweezer at room temperature. After 5 minutes of incubation, the sample was manually blotted away by gently touching the rim of the grid with a Whatman 1 filter paper (Whatman, CAT No.1001-090) to wick away the unbound protein solution and replaced with another 3.5 µL. This was repeated for a total of 4-6 times. To get rid of the sucrose in γ-TuRC storage buffer containing ∼30% sucrose and to pre-equilibrate γ-TuRC with microtubule nucleation compatible buffer, 20 µL of assay buffer (1X BRB80 (80 mM PIPES, 1 mM MgCl_2_, 1 mM EGTA), 1 mM GTP, 50 mM KCl, 1 mM DTT, pH 6.8, adjusted with KOH) was applied to the grid and wicked away with a Whatman 1 filter paper after 5 minutes of incubation. This was followed by a second wash step with 5 µL of assay buffer application and the vitrobot tweezer with the grid was then moved to a heat block at 37°C and covered with a lid to make a closed chamber. After 2 minutes of incubation, the buffer was carefully blotted away with a filter paper and replaced with a 5 µL nucleation mixture containing 10 µM E254D tubulin and 100 nM chTOG in the assay buffer. The nucleation/polymerization reaction was allowed to proceed for 50 s on the heat block at 37°C and then the tweezer was transferred to the Vitrobot IV (Thermo Scientific) arm, pre-equilibrated to 37°C. The grid was blotted for 3-3.5 s at 100% humidity and 37°C and then plunge frozen in liquid ethane.

Data was collected in 7 different sessions on FEI Titan Krios microscopes using Serial EM automated data collection (*45*) for the 1st dataset and using leginon (*46*) for the rest of the 6 datasets (2–7). The data acquisition parameters are summarized in the table S3.

#### TIRF-based single molecule γ-TuRC-GFP microtubule nucleation assays

Microtubule nucleation assays were performed as described in (*24*) on a Nikon Eclipse Ti TIRF microscope setup equipped with a NA-1.49 100X Plan Apo TIRF objective. It also included a 3-axis piezo-electric stage (Mad City LabsNano LP-200) and two-color TIRF imaging optics (lasers: 488 nm (Spectra-physics) and 561 nm (Cobalt); Filters: Emission (Semrock FF01-520/35 and FF01-609/54), Dichroic (Semrock Di01-R488/561). Images were acquired using a Photometrics Prime 95B sCMOS camera. Glass slides (TED PELLA, 260600) and glass coverslips (Fisherbrand, 12542A) were cleaned using the following steps: i) Rinsing in acetone for 5 minutes, ii) Sonication in 50% (volume/volume) Methanol in milli-Q water for 20 minutes in a Branson Ultrasonics water bath, iii) Sonication in 0.5 M KOH for 20 minutes in a Branson Ultrasonics water bath, iv) Rinsing in milli-Q water followed by drying with nitrogen gas. The glass coverslips were then air-plasma cleaned for 10 minutes (Harrick Plasma PDC-32G). Flow cells were assembled using 2 strips of double-sided tapes (Scotch) on a glass slide and adhering a cleaned coverslip onto the tape resulting in a flow cell of 8-10 µL in volume. The flow cell was pre-rinsed with 1XBRB80 - 1 mM tris (2 carboxyethyl) phosphine (TCEP) followed by incubation with 0.2 mg/mL poly-L-lysine (PLL) - polyethylene glycol (PEG) - biotin (PLL[20]- g [3.5] - PEG[2] /PEG[3.4] - biotin [20%]; SUSOS AG) prepared in 1XBRB80 + 1 mM TCEP for 5 minutes in a moist closed chamber at room temperature. The flow cell was then washed with 1XBRB80 - 1 mM TCEP followed by addition of a mixture of 0.5 mg/mL κ-casein (Millipore Sigma) and 0.25 mg/mL neutravidin in 1XBRB80 - 1 mM TCEP (Thermo FisherScientific). After 5 minutes of incubation, the flow cell was washed with assay buffer (1XBRB80 - 1 mM TCEP, 50 mM KCl, 0.15% [weight/volume] methylcellulose, 0.2 mg/mL κ-casein and 1 mM GTP). This was followed by flow-in of 0.02 mg/mL biotinylated GFP nanobody diluted in the assay buffer. The unbound GFP nanobody was washed away after 5 minutes by flowing in assay buffer in the flow cell. γ-TuRC, freshly diluted to ∼1 pM in assay buffer, was then flowed in the flow cell and incubated for 5 minutes. Following a wash with assay buffer, the nucleation/polymerization mixture containing 10 µM E254D tubulin or wild type tubulin mixed with 11 mol% X-rhodamine bovine brain tubulin, 100 nM chTOG and oxygen scavenging mixture (0.035 mg/mL catalase, 0.2 mg/mL glucose oxidase, 2.5 mM glucose, and 10 mM DTT) in assay buffer was introduced in the flow cell. The open ends of the flow cell were sealed with VALAP (1:1:1 petroleum jelly/ lanolin/ paraffin) and the assay was imaged on the TIRF microscope stage, preheated to 37°C. Time lapse images were acquired in the GFP (green) and tubulin (red) channels sequentially with an exposure time of 300 ms and 500 ms respectively for the two channels separated by 5 s-intervals for a total time of 5 minutes. Images were acquired using NIS-Elements AR 4.60.00 (Nikon).The time-lapse images were analyzed in Fiji and γ-TuRC-mediated nucleation events were counted and tracked manually for the wild-type and E254D tubulin. Nucleation events initiating from a green fluorescent puncta and showing microtubule growth only at the other non-capped end were classified as γ-TuRC-GFP foci nucleated, whereas nucleation events showing microtubule growth at both the ends were classified as spontaneous nucleation events in fig. S1D-F. For fig. S1G, only nucleation events initiating from a green fluorescent puncta were considered.

### 3D classification for protofilament sorting

#### γ-TuRC nucleated microtubules

177 microtubules with a cone-shaped density at the end of a microtubule were manually marked in multiple rounds of careful inspection of micrographs from 5 different grids (table S1). Microtubule segments were classified into different protofilament numbers based on the Microtubule Relion Pipeline (MiRP) (*47*). Microtubule start and end coordinates were manually picked for these microtubules in RELION for straight, non-overlapping segments and particles were extracted by binning them 4X, starting with a box size of 512 pixels (1.32 Å/pixel). The first 6 segments starting from the cone-shaped density were removed to exclude γ-TuRC particles from analysis. Only microtubules that had at least 5 segments were analyzed (150 out of 177 microtubules). Binned segments were subjected to 2D classification, followed by a single round of 3D classification with alignment to synthetic references of microtubules of different protofilament architecture (11-16 protofilaments). A microtubule was assigned a class/protofilament number if all of the 5 consecutive particles starting from the minus end side of that microtubule belonged to the same 3D class. Microtubules with particles belonging to more than one class in the first 5 consecutive particles starting from the minus end side of the microtubule were not assigned a class (25 out of 150 microtubules).

#### Spontaneously nucleated microtubules

For spontaneously nucleated microtubules, microtubule start and end coordinates were manually picked for microtubules in RELION for straight, non-overlapping segments and particles were extracted by binning them 4X, starting with a box size of 512 pixels (1.32 Å/pixel). Microtubules with at least 5 segments were analyzed as above (225 out of 228 microtubules). A microtubule was assigned a class/protofilament number if a minimum of 5 consecutive particles in the microtubule segment belonged to that class. Microtubules with two or more 5 consecutive particle stretches belonging to 2 different 3D classes were not assigned a class (19 out of 225 microtubules).

### Cryo-EM data processing

#### γ-TuRC-capped microtubule minus-end

Inter-frame local motion correction and dose-weighting were performed using MotionCor2 (*48*). Contrast transfer function (CTF) parameters for the drift-corrected and dose-weighted micrographs were estimated using CTFFIND4 (*49*). RELION v.3.1 (*50*) was used for subsequent processing unless noted otherwise. Dose-weighted micrographs were manually inspected, and microtubule ends with a cone-shaped density were picked. These microtubule ends were binned to a pixel size of 5.28 Å/pixel and extracted with a box size of 180 pixels. Multiple rounds of reference-free 2D classification and re-extraction with the ‘re-center’ option selected, were performed to select a subset of particles with clear spoke-like densities at the microtubule end and align them in the middle of the box. These cleaned particles were used to generate an *ab initio* model. Particles were re-extracted with a pixel size of 1.32 Å/pixel and box size of 360 pixels. Re-extracted particles were subjected to a round of 3D refinement using the *ab initio* model from above. Signal outside the stable region of the map, which includes positions 1-2 and 10-14, was subtracted from the raw particles using the ‘Particle subtraction’ function in RELION. Signal-subtracted particles were subjected to focused 3D refinement yielding a map with sufficient resolution for CTF refinement (per-micrograph beam tilt, B-factor, astigmatism, defocus, and anisotropic magnification). Subsequently, the ‘revert to original particles’ option was used to transfer the refined CTF parameters to the raw particles used for local 3D refinement, yielding a 7.7Å resolution map of the γ-TuRC-capped microtubule minus-end.

To improve the density at the dynamic region of the structure, we subtracted densities corresponding to α/β-tubulin dimers from the raw particle images using the ‘Particle subtraction’ function. Subtracted particles were then subjected to multiple rounds of 3D classification without alignments to obtain a subset of particles with the strongest signal for spoke densities in the dynamic region. The final subset of 8,195 particles were reverted to the original raw particle images and subjected to local 3D refinement, yielding an overall resolution of 9.5Å.

A similar strategy using particle subtraction and focused 3D refinement was implemented to improve the reconstruction at the stable region of the structure corresponding to positions 3 to 9, yielding a 7.5Å resolution map. Coordinates for γ-tubulin (AlphaFold: AF-P23258-F1), and α-tubulin (PDB ID: 6DPV) (*51*) were individually rigid-body docked into positions 3-9. As pairwise comparison of adjacent tubulins revealed no significant structural difference, symmetry-expansion was performed based on the helical parameters: helical rise = 9.46 Å and helical twist = 27.69°. Particle subtraction with a mask covering four tubulins (a pair of γ-tubulins and the proximal pair of α-tubulins contacting them) was performed, followed by focused 3D refinement and 3D classification with no alignments. Particles corresponding to the best class were subjected to local 3D refinement, yielding a 4.0Å map.

#### Free recombinant γ-TuRC

The structure of the free recombinant γ-TuRC was determined using a previously described strategy (*8*). Briefly, a small set of free γ-TuRC particles were manually selected from a few micrographs and subjected to reference-free 2D classification. The 2D class averages generated were used as references to autopick particles from all micrographs. The autopicked particles were extracted and binned to the pixel size of 10.93 Å/pixel. Multiple rounds of reference-free 2D classification were used to select free γ-TuRC particles. These particles were then subjected to one round of 3D refinement followed by re-extraction with the ‘re-center’ option selected and a pixel size of 5.32 Å/pixel. This was followed by another round of 3D refinement and 3D classification to select “good” γ-TuRC particles. Particles corresponding to the best 3D classes were re-extracted with a pixel size of 2 Å/pixel and subjected to an additional round of 2D classification to select 933,254 clean particles. These clean particles were imported into CryoSPARC v.4.0.2 (*52*) and subjected to heterogeneous refinement. A subset of 152,604 particles corresponding to the 3D classes with the best density throughout the γ-TuRC were selected and used to run non-uniform refinement, yielding an overall density map at 7.3Å.

#### Model building

Coordinates for individual GCPs were extracted from the native human γ-TuRC structure (PDB ID: 6v6s) (*8*). The coordinates were further segmented into two parts (N- and C-domains for each GCP). N- and C-domains of GCP-2 were individually rigid-body fitted into the GCP density at position 7 of the “positions 3 to 9” map since the densities for GCP and γ-tubulin were at the highest resolution in this position (fig. S3). The alpha-fold model (AF-P23258-F1) for the full-length γ-tubulin was used for rigid-body fitting into the γ-tubulin density at position 7. This GCP-γ-tubulin module was rigid body fitted into the rest of the spokes and showed good agreement with the EM density.

Coordinates for the GCP N-domains at positions 1-12 from PDB ID: 6v6s (*8*) were fitted as a unit into the “dynamic” γ-TuRC-capped microtubule minus-end map. The C-domain of GCP and γ-tubulin in position 7 from the “positions 3 to 9” map was fit into corresponding densities for GCP and γ-tubulin in each spoke in positions 3-12. Coordinates for the hetero-tetramer containing GCP-2, GCP-3 and 2 copies of γ-tubulin from positions 7-8 from the “positions 3 to 9” map were rigid-body fitted into positions 1-2 and 13-14. The separately fitted components were combined to generate a consensus model for free and microtubule-bound γ-TuRCs. Coordinates for α/β-tubulin (PDB ID: 6DPV) (*51*) were rigid body fitted into the tubulin densities in the “dynamic” map to generate the γ-TuRC-capped microtubule minus-end model. α-tubulin from α/β-tubulin coordinates (PDB ID: 6DPV) (*51*) were used to rigid body fit into the densities corresponding to α-tubulins in the symmetry expanded map of adjacent γ-tubulin-α-tubulin pairs (fig. S3), which was used to generate a model for tubulin lateral interaction interfaces.

Rigid-body fitting into cryo-EM densities was performed in UCSF Chimera, a, developed by the Resource for Biocomputing, Visualization, and Informatics at the University of California, San Francisco, with support from NIH P41-GM103311 and inspected in COOT (*53*). Figures were generated in PyMol (The PyMOL Molecular Graphics System, Version 2.0 Schrödinger, LLC.).

## Acknowledgement

This work was funded by an NIH grant (GM130234) to TMK. AA was supported by a Pels Family Center Postdoctoral Fellowship at The Rockefeller University. Some of this work was performed at the Simons Electron Microscopy Center (SEMC) at the New York Structural Biology Center, with major support from the Simons Foundation (SF349247). The authors are grateful to Dr. Joshua Mendez, SEMC for help with data acquisition. The authors are grateful to M. Ebrahim, J. Sotiris, and H.Ng and the Evelyn Gruss Lipper Cryo-Electron Microscopy Resource Center for cryo-EM support and Dr. Hilda Amalia Pasolli at the Electron Microscopy Resource Center, both at Rockefeller University for EM support.

## Author contributions

Conceptualization: AA, LU and TMK

Methodology: AA and LU

Investigation: AA, LU, AX and KN

Visualization: AA, LU and TMK

Funding acquisition: TMK

Project administration: TMK

Supervision: TMK

Writing – original draft: AA, LU and TMK

Writing – review & editing: AA, LU and TMK

## REFERENCES

1. M. Kirschner, T. Mitchison, Beyond self-assembly: from microtubules to morphogenesis. Cell. 45, 329–342 (1986).

2. G. J. Brouhard, L. M. Rice, Microtubule dynamics: an interplay of biochemistry and mechanics. Nat. Rev. Mol. Cell Biol. 19, 451–463 (2018).

3. A. Akhmanova, M. O. Steinmetz, Control of microtubule organization and dynamics: two ends in the limelight. Nat. Rev. Mol. Cell Biol. 16, 711–726 (2015).

4. L. G. Tilney, J. Bryan, D. J. Bush, K. Fujiwara, M. S. Mooseker, D. B. Murphy, D. H. Snyder, Microtubules: evidence for 13 protofilaments. J. Cell Biol. 59, 267–275 (1973).

5. K. J. Böhm, W. Vater, H. Fenske, E. Unger, Effect of microtubule-associated proteins on the protofilament number of microtubules assembled in vitro. Biochim. Biophys. Acta. 800, 119–126 (1984).

6. L. Evans, T. Mitchison, M. Kirschner, Influence of the centrosome on the structure of nucleated microtubules. J. Cell Biol. 100, 1185–1191 (1985).

7. D. Chrétien, F. Metoz, F. Verde, E. Karsenti, R. H. Wade, Lattice defects in microtubules: protofilament numbers vary within individual microtubules. J. Cell Biol. 117, 1031–1040 (1992).

8. M. Wieczorek, L. Urnavicius, S.-C. Ti, K. R. Molloy, B. T. Chait, T. M. Kapoor, Asymmetric Molecular Architecture of the Human γ-Tubulin Ring Complex. Cell. 180, 165–175.e16 (2020).

9. P. Liu, E. Zupa, A. Neuner, A. Böhler, J. Loerke, D. Flemming, T. Ruppert, T. Rudack, C. Peter, C. Spahn, O. J. Gruss, S. Pfeffer, E. Schiebel, Insights into the assembly and activation of the microtubule nucleator γ-TuRC. Nature. 578, 467–471 (2020).

10. T. Consolati, J. Locke, J. Roostalu, Z. A. Chen, J. Gannon, J. Asthana, W. M. Lim, F. Martino, M. A. Cvetkovic, J. Rappsilber, A. Costa, T. Surrey, Microtubule Nucleation Properties of Single Human γTuRCs Explained by Their Cryo-EM Structure. Dev. Cell. 53, 603–617.e8 (2020).

11. F. Zimmermann, M. Serna, A. Ezquerra, R. Fernandez-Leiro, O. Llorca, J. Luders, Assembly of the asymmetric human γ-tubulin ring complex by RUVBL1-RUVBL2 AAA ATPase. Science Advances. 6, eabe0894 (2020).

12. M. Wieczorek, T.-L. Huang, L. Urnavicius, K.-C. Hsia, T. M. Kapoor, MZT Proteins Form Multi-Faceted Structural Modules in the γ-Tubulin Ring Complex. Cell Rep. 31, 107791 (2020).

13. Y. Zheng, M. L. Wong, B. Alberts, T. Mitchison, Nucleation of microtubule assembly by a gamma-tubulin-containing ring complex. Nature. 378, 578–583 (1995).

14. M. Moritz, M. B. Braunfeld, V. Guénebaut, J. Heuser, D. A. Agard, Structure of the γ-tubulin ring complex: a template for microtubule nucleation. Nat. Cell Biol. 2, 365–370 (2000).

15. T. J. Keating, G. G. Borisy, Immunostructural evidence for the template mechanism of microtubule nucleation. Nat. Cell Biol. 2, 352–357 (2000).

16. C. Wiese, Y. Zheng, A new function for the gamma-tubulin ring complex as a microtubule minus-end cap. Nat. Cell Biol. 2, 358–364 (2000).

17. J. M. Kollman, C. H. Greenberg, S. Li, M. Moritz, A. Zelter, K. K. Fong, J.-J. Fernandez, A. Sali, J. Kilmartin, T. N. Davis, D. A. Agard, Ring closure activates yeast γTuRC for species-specific microtubule nucleation. Nat. Struct. Mol. Biol. 22, 132–137 (2015).

18. J. M. Kollman, A. Merdes, L. Mourey, D. A. Agard, Microtubule nucleation by γ-tubulin complexes. Nat. Rev. Mol. Cell Biol. 12, 709–721 (2011).

19. B. R. Oakley, V. Paolillo, Y. Zheng, γ-Tubulin complexes in microtubule nucleation and beyond. Mol. Biol. Cell. 26, 2957–2962 (2015).

20. M. Moritz, Y. Zheng, B. M. Alberts, K. Oegema, Recruitment of the gamma-tubulin ring complex to Drosophila salt-stripped centrosome scaffolds. J. Cell Biol. 142, 775–786 (1998).

21. K. Oegema, C. Wiese, O. C. Martin, R. A. Milligan, A. Iwamatsu, T. J. Mitchison, Y. Zheng, Characterization of Two Related Drosophila γ-tubulin Complexes that Differ in Their Ability to Nucleate Microtubules. J. Cell Biol. 144, 721–733 (1999).

22. J. M. Kollman, A. Zelter, E. G. D. Muller, B. Fox, L. M. Rice, T. N. Davis, D. A. Agard, The Structure of the γ-Tubulin Small Complex: Implications of Its Architecture and Flexibility for Microtubule Nucleation. MBoC. 19, 207–215 (2008).

23. J. M. Kollman, J. K. Polka, A. Zelter, T. N. Davis, D. A. Agard, Microtubule nucleating gamma-TuSC assembles structures with 13-fold microtubule-like symmetry. Nature. 466, 879–882 (2010).

24. M. Wieczorek, S.-C. Ti, L. Urnavicius, K. R. Molloy, A. Aher, B. T. Chait, T. M. Kapoor, Biochemical reconstitutions reveal principles of human γ-TuRC assembly and function (2020), p. 2020.09.21.306845.

25. A. Y. Berman, M. Wieczorek, A. Aher, P. D. B. Olinares, B. T. Chait, T. M. Kapoor, A nucleotide binding–independent role for γ-tubulin in microtubule capping and cell division. J. Cell Biol. 222 (2023), doi:10.1083/jcb.202204102.

26. G. J. Brouhard, J. H. Stear, T. L. Noetzel, J. Al-Bassam, K. Kinoshita, S. C. Harrison, J. Howard, A. A. Hyman, XMAP215 is a processive microtubule polymerase. Cell. 132, 79– 88 (2008).

27. A. Thawani, R. S. Kadzik, S. Petry, XMAP215 is a microtubule nucleation factor that functions synergistically with the γ-tubulin ring complex. Nat. Cell Biol. 20, 575–585 (2018).

28. C. Guyomar, C. Bousquet, S. Ku, J. M. Heumann, G. Guilloux, N. Gaillard, C. Heichette, L. Duchesne, M. O. Steinmetz, R. Gibeaux, D. Chrétien, Changes in seam number and location induce holes within microtubules assembled from porcine brain tubulin and in Xenopus egg cytoplasmic extracts. Elife. 11 (2022), doi:10.7554/eLife.83021.

29. G. B. Pierson, P. R. Burton, R. H. Himes, Alterations in number of protofilaments in microtubules assembled in vitro. J. Cell Biol. 76, 223–228 (1978).

30. E. Unger, K. J. Böhm, W. Vater, Structural diversity and dynamics of microtubules and polymorphic tubulin assemblies. Electron Microsc. Rev. 3, 355–395 (1990).

31. J. Roostalu, C. Thomas, N. I. Cade, S. Kunzelmann, I. A. Taylor, T. Surrey, The speed of GTP hydrolysis determines GTP cap size and controls microtubule stability. Elife. 9 (2020), doi:10.7554/eLife.51992.

32. G. M. Alushin, G. C. Lander, E. H. Kellogg, R. Zhang, D. Baker, E. Nogales, High-resolution microtubule structures reveal the structural transitions in αβ-tubulin upon GTP hydrolysis. Cell. 157, 1117–1129 (2014).

33. R. Kiewisz, G. Fabig, W. Conway, D. Baum, D. Needleman, T. Müller-Reichert, Three-dimensional structure of kinetochore-fibers in human mitotic spindles. Elife. 11 (2022), doi:10.7554/eLife.75459.

34. P. Guichard, D. Chrétien, S. Marco, A.-M. Tassin, Procentriole assembly revealed by cryo-electron tomography. EMBO J. 29, 1565–1572 (2010).

35. M. C. Pamula, L. Carlini, S. Forth, P. Verma, S. Suresh, W. R. Legant, A. Khodjakov, E. Betzig, T. M. Kapoor, High-resolution imaging reveals how the spindle midzone impacts chromosome movement. J. Cell Biol. 218, 2529–2544 (2019).

36. E. Nogales, R. Zhang, Visualizing microtubule structural transitions and interactions with associated proteins. Curr. Opin. Struct. Biol. 37, 90–96 (2016).

37. S. W. Manka, C. A. Moores, Microtubule structure by cryo-EM: snapshots of dynamic instability. Essays Biochem. 62, 737–751 (2018).

38. M. Würtz, E. Zupa, E. S. Atorino, A. Neuner, A. Böhler, A. S. Rahadian, B. J. A. Vermeulen, G. Tonon, S. Eustermann, E. Schiebel, S. Pfeffer, Modular assembly of the principal microtubule nucleator γ-TuRC. Nat. Commun. 13, 473 (2022).

39. S.-C. Ti, G. M. Alushin, T. M. Kapoor, Human β-tubulin isotypes can regulate microtubule protofilament number and stability. Dev. Cell. 47, 175–190.e5 (2018).

40. M. P. Miller, C. L. Asbury, S. Biggins, A TOG Protein Confers Tension Sensitivity to Kinetochore-Microtubule Attachments. Cell. 165, 1428–1439 (2016).

41. Y.-K. Choi, P. Liu, S. K. Sze, C. Dai, R. Z. Qi, CDK5RAP2 stimulates microtubule nucleation by the γ-tubulin ring complex. J. Cell Biol. 191, 1089–1095 (2010).

42. P. C. Fridy, Y. Li, S. Keegan, M. K. Thompson, I. Nudelman, J. F. Scheid, M. Oeffinger, M. C. Nussenzweig, D. Fenyö, B. T. Chait, M. P. Rout, A robust pipeline for rapid production of versatile nanobody repertoires. Nat. Methods. 11, 1253–1260 (2014).

43. S. M. Abmayr, T. Yao, T. Parmely, J. L. Workman, Curr. Protoc. Mol. Biol., in press, doi:10.1002/0471142727.mb1201s75.

44. J. Snijder, A. J. Borst, A. Dosey, A. C. Walls, A. Burrell, V. S. Reddy, J. M. Kollman, D. Veesler, Vitrification after multiple rounds of sample application and blotting improves particle density on cryo-electron microscopy grids. J. Struct. Biol. 198, 38–42 (2017).

45. D. N. Mastronarde, Automated electron microscope tomography using robust prediction of specimen movements. J. Struct. Biol. 152, 36–51 (2005).

46. C. Suloway, J. Pulokas, D. Fellmann, A. Cheng, F. Guerra, J. Quispe, S. Stagg, C. S. Potter, B. Carragher, Automated molecular microscopy: The new Leginon system. J. Struct. Biol. 151, 41–60 (2005).

47. A. D. Cook, S. W. Manka, S. Wang, C. A. Moores, J. Atherton, A microtubule RELION-based pipeline for cryo-EM image processing. J. Struct. Biol. 209, 107402 (2020).

48. S. Q. Zheng, E. Palovcak, J.-P. Armache, K. A. Verba, Y. Cheng, D. A. Agard, MotionCor2: anisotropic correction of beam-induced motion for improved cryo-electron microscopy. Nat. Methods. 14, 331–332 (2017).

49. A. Rohou, N. Grigorieff, CTFFIND4: Fast and accurate defocus estimation from electron micrographs. J. Struct. Biol. 192, 216–221 (2015).

50. J. Zivanov, T. Nakane, B. O. Forsberg, D. Kimanius, W. J. Hagen, E. Lindahl, S. H. Scheres, New tools for automated high-resolution cryo-EM structure determination in RELION-3. Elife. 7 (2018), doi:10.7554/eLife.42166.

51. R. Zhang, B. LaFrance, E. Nogales, Separating the effects of nucleotide and EB binding on microtubule structure. Proc. Natl. Acad. Sci. U. S. A. 115, E6191–E6200 (2018).

52. A. Punjani, J. L. Rubinstein, D. J. Fleet, M. A. Brubaker, cryoSPARC: algorithms for rapid unsupervised cryo-EM structure determination. Nat. Methods. 14, 290–296 (2017).

53. P. Emsley, B. Lohkamp, W. G. Scott, K. Cowtan, Features and development of Coot. Acta Crystallogr. D Biol. Crystallogr. 66, 486–501 (2010).

