## Supplementary figures for "Structure of the γ-tubulin ring complex-capped microtubule"

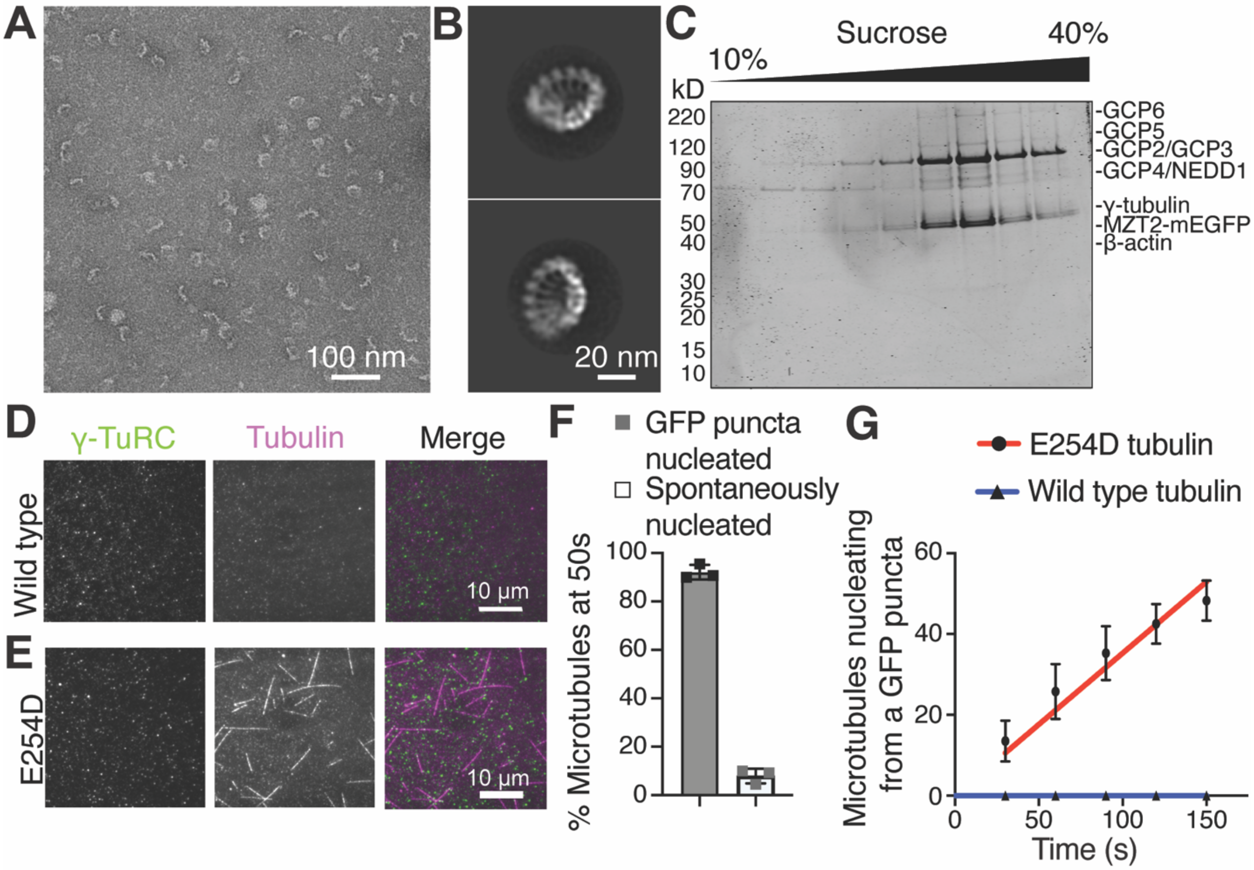


**Fig. S1. TIRF-based optimization of γ-TuRC-dependent microtubule nucleation assay with slow GTP hydrolysing mutant of α-tubulin, E254D. (A)** Transmission EM micrograph of negatively stained native γ-TuRC. **(B)** 2D class-averages showing two orientations of native γ-TuRC particles. **(C)** Coomassie stained SDS-PAGE analysis of recombinant γ-TuRC-GFP after sucrose density gradient centrifugation. **(D,E)** Stills from a TIRF-movie showing the γ-TuRC (green), tubulin (magenta) and merge channels after 2 minutes 30 s in the presence of 100 nM chTOG and either 10 μM wild type tubulin (D) or 10 μM E254D (TUBA1B-E254D, TUBB3) tubulin (E). **(F)** Plot of the percentage of microtubules nucleated by a γ-TuRC-GFP puncta or spontaneously at 50s post start of nucleation in the presence of 10 μM E254D tubulin and 100 nM chTOG. **(G)** Plot of the cumulative number of microtubules nucleated by γ-TuRC in the presence of 100 nM chTOG and either 10 μM wild type tubulin or 10 μM E254D tubulin over time. Data were fitted using linear regression, red line for E254D tubulin and blue line for wild type tubulin. n=2 replicates for wild type tubulin and n=4 replicates for E254D tubulin.

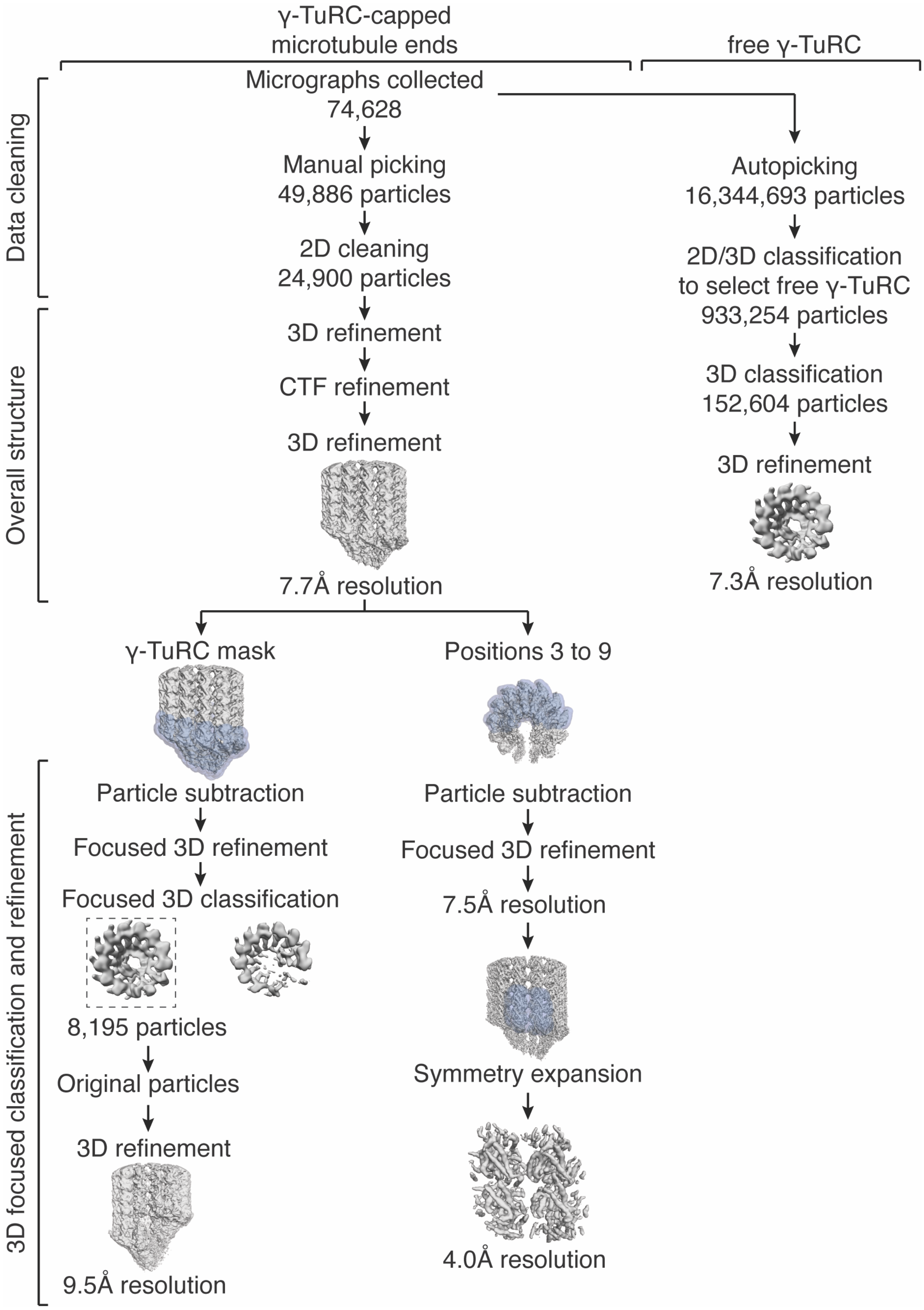
γ-TuRC-capped microtubule ends
free γ-TuRC
Data cleaning
Overall structure
3D focused classification and refinement
Micrographs collected
74,628
Manual picking
49,886 particles
2D cleaning
24,900 particles
3D refinement
CTF refinement
3D refinement
7.7Å resolution
Autopicking
16,344,693 particles
2D/3D classification
to select free γ-TuRC
933,254 particles
3D classification
152,604 particles
3D refinement
7.3Å resolution
γ-TuRC mask
Positions 3 to 9
Particle subtraction
Focused 3D refinement
Focused 3D classification
8,195 particles
Original particles
3D refinement
9.5Å resolution
Particle subtraction
Focused 3D refinement
7.5Å resolution
Symmetry expansion
4.0Å resolution

**Fig. S2. Cryo-EM data processing workflow for γ-TuRC-capped microtubule minus-end and free γ-TuRC.**

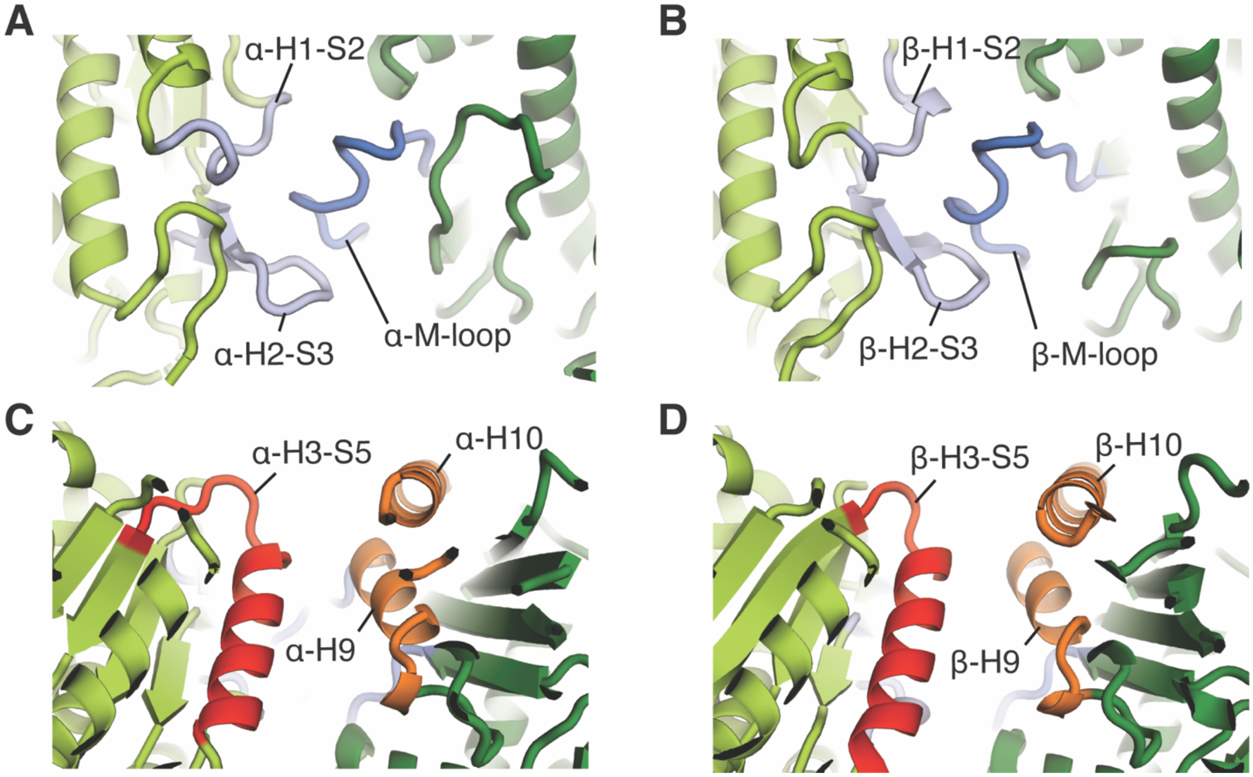


**Fig. S3. Lateral α-α and β-β tubulin interactions.** Lateral interactions involving the M-loop at α-tubulin-α-tubulin and β-tubulin-β-tubulin interface (ribbons) respectively **(A,B)** and the H3,H9 and H10 helices **(C,D)**.

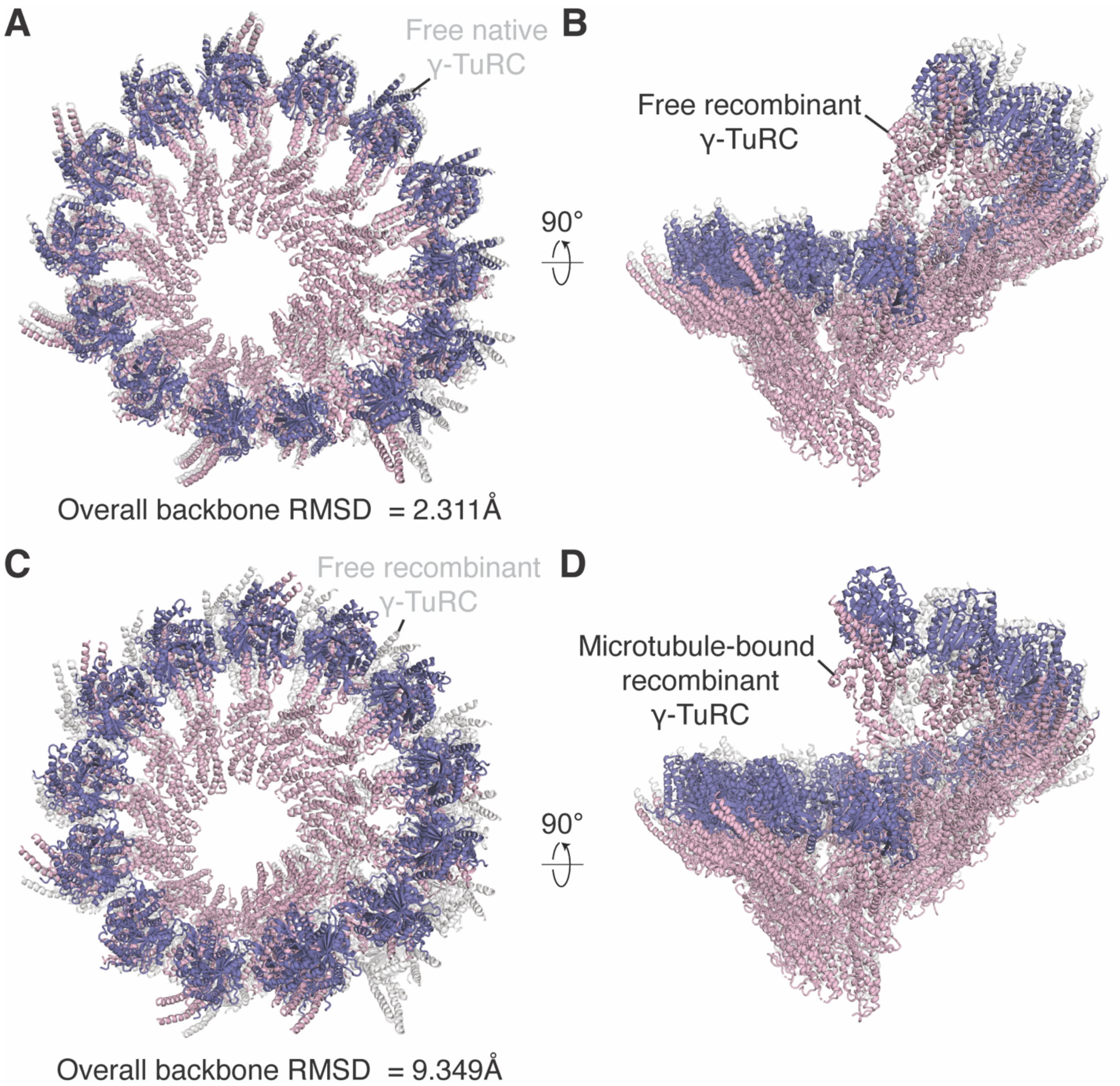


**Fig. S4. Comparison of γ-TuRC in its free and microtubule bound form.** (**A,B**) Comparison of the rigid-body fitted models of native γ-TuRC and recombinant γ-TuRC showing top (A) and side views (B). **(C,D)** Comparison of the rigid-body fitted models of recombinant γ-TuRC and microtubule bound recombinant γ-TuRC showing top (C) and side views (D).

| Dataset | 1 | 2 | 3 | 4 | 5 |
|---|---|---|---|---|---|
| Acceleration voltage (kV) | 300 | 300 | 300 | 300 | 300 |
| Camera | K2 | K3 | K3 | K3 | K2 |
| Super-resolution mode | Yes | Yes | Yes | Yes | Yes |
| CS corrector | No | Yes | Yes | Yes | No |
| Energy filter (20 $e^-$-slit) | No | Yes | Yes | Yes | No |
| Exposure time (s) | 8 | 3 | 3 | 3 | 8 |
| Frame number | 20 | 20 | 20 | 20 | 20 |
| Pixel size (Å/pixel) | 1.32 | 1.32 | 1.32 | 1.32 | 1.32 |
| Dose rate ($e^-$/Å$^2$/s) | 4.49 | 11.48 | 11.48 | 11.48 | 4.49 |
| Total dose ($e^-$/Å$^2$) | 35.91 | 34.44 | 34.44 | 34.44 | 35.91 |

**Table S1. Cryo-EM data collection parameters for native γ-TuRC nucleated microtubules.**

| Dataset | 1 |
|---|---|
| Acceleration voltage (kV) | 300 |
| Camera | K2 |
| Super-resolution mode | Yes |
| CS corrector | No |
| Energy filter (20 $e^-$-slit) | No |
| Exposure time (s) | 8 |
| Frame number | 20 |
| Pixel size (Å/pixel) | 1.32 |
| Dose rate ($e^-$/Å$^2$/s) | 4.49 |
| Total dose ($e^-$/Å$^2$) | 35.91 |

**Table S2. Cryo-EM data collection parameters for spontaneously nucleated microtubules.**

| Dataset | 1 | 2 | 3 | 4 | 5 | 6 | 7 |
|---|---|---|---|---|---|---|---|
| Acceleration voltage (kV) | 300 | 300 | 300 | 300 | 300 | 300 | 300 |
| Camera | K3 | K3 | K3 | K3 | K3 | K3 | K3 |
| Super-resolution mode | Yes | Yes | Yes | Yes | Yes | Yes | Yes |
| CS corrector | Yes | Yes | No | No | No | Yes | No |
| Energy filter (20 $e^-$-slit) | Yes | Yes | Yes | Yes | Yes | Yes | Yes |
| Exposure time (s) | 4 | 2.3 | 2.8 | 2.3 | 2.3 | 2.0 | 2.4 |
| Frame number | 40 | 46 | 56 | 58 | 58 | 40 | 48 |
| Pixel size (Å/pixel) | 1.32 | 1.076 | 1.083 | 1.083 | 1.083 | 1.076 | 1.083 |
| Dose rate ($e^-$/Å$^2$/s) | 14.35 | 25.93 | 21.31 | 25.64 | 25.64 | 26.06 | 21.81 |
| Total dose ($e^-$/Å$^2$) | 57.39 | 59.64 | 59.66 | 58.97 | 58.98 | 52.12 | 52.35 |

**Table S3. Cryo-EM data collection parameters for recombinant γ-TuRC nucleated microtubules.**
